# Targeting HMGA1-driven leukemic transformation in myeloproliferative neoplasms with pacritinib

**DOI:** 10.1101/2025.06.01.657170

**Authors:** Can Yang, Yajun Li, Kun Chen, Xuemei Tang, Qingyun Zhang, Wenjiao Qin, Yu Zeng, Jianfei Fu, Xiaotong Ma, Wenshu Zheng, Zhen Huang, Wenhao Zhang, Gusheng Tang, Ming Guan, Zhiyuan Wu

## Abstract

Progression of myeloproliferative neoplasms (MPNs) to secondary acute myeloid leukemia (sAML) is a lethal transition lacking reliable early biomarkers and effective therapies. Integrating multi-omics analyses, a cohort of 240 patient biopsies, and functional studies, we identify High Mobility Group A1 (HMGA1) as a critical driver and predictor of this transformation. HMGA1 expression, readily detectable by immunohistochemistry, progressively increases from chronic MPN phases to sAML, outperforming established markers (CD34/CD117) in predicting impending leukemic transformation (AUC = 0.96). Mechanistically, HMGA1 cooperates with *JAK2*^V617F^ and *TP53* mutations to promote leukemogenesis. Clinically, high HMGA1 levels correlate with resistance to first-generation JAK inhibitors and portend poor overall survival. Importantly, the next generation JAK2/FLT3 inhibitor pacritinib abrogates HMGA1-driven proliferation and significantly extends survival in preclinical models, offering an immediate therapeutic strategy for this high-risk population. Our findings establish HMGA1 as an actionable biomarker for risk stratification and early intervention, and a therapeutic target to overcome therapeutic resistance in MPN-sAML.

**Highlights:**

1. HMGA1 IHC staining serves as a readily implementable biomarker, outperforming existing marker for early prediction of MPN progression to sAML and identifying patients at high risk.
2. Elevated HMGA1 expression correlates with resistance to first-generation JAK inhibitors and predicts poor overall survival in MPN-sAML patients, highlighting a critical unmet therapeutic need.
3. The next-generation JAK2 inhibitor pacritinib effectively overcomes HMGA1-driven aggressive disease and therapy resistance, offering a rational, clinically available treatment strategy for HMGA1-high MPN-sAML.

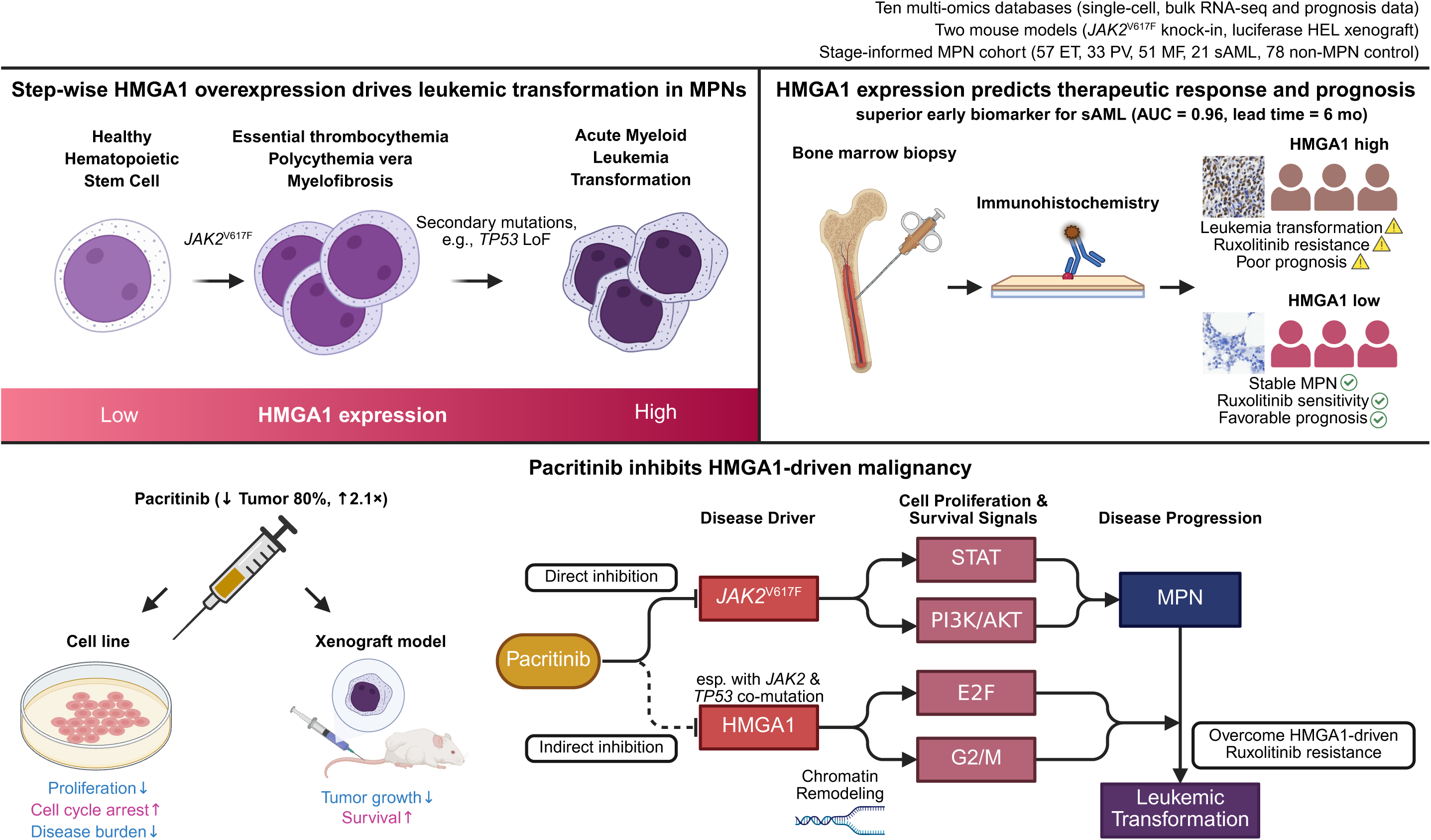

## 1 Introduction

The transformation of Philadelphia-negative myeloproliferative neoplasms (MPNs) into secondary acute myeloid leukemia (sAML) represents a critical and often fatal juncture in disease progression, presenting formidable challenges in both early diagnosis and effective therapeutic intervention due to sAML’s adverse-risk nature and frequent treatment resistance^1^. With a median survival under six months^2^, this aggressive leukemia can develop in up to 20% of MPNs patients, who are primarily driven by mutations such as *JAK2*^V617F^, *CALR* exon-9 indels or *MPL*^W515 3,4^. Leukemic progression involves significant genetic complexity, dynamic cloncal changes, and sequential acquisition of high-risk genetic lesions^5^. Notably, *TP53* mutations are prominent in 25-50% of post-MPN sAML, often driving clonal expansion^6^ and alter therapy responses^7^. The acquisition of further oncogenic mutations, including those in epigenetic modifiers and splicing factors, also marks this progression^8–10^.

A paramount challenge in MPN management is the limited capacity of established clinical risk stratification models (IPSS, DIPSS, MIPSS-70, ELN) to accurately predict sAML transformation and inform pre-emptive therapeutic strategies. These scoring systems often fail to fully encapsulate the dynamic clonal evolution driving disease progression^1,11^, delaying critical interventions like allogeneic stem cell transplantation until overt sAML (blast burden ≥10 or 20%), when clinical outcomes are profoundly compromised^12,13^. Moreover, first-generation JAK2 inhibitors like ruxolitinib, while palliating chronic-phase MPN, confer marginal benefit in sAML and may inadvertently select for drug-resistant subclones^14^, partly due to underlying molecular mechanisms of persistence such as aberrant signaling pathway activity^15^. Consequently, a pressing unmet clinical need persists for robust, early biomarkers to identify high-rsik patients and for innovative therapeutic modalities to intercept or effectively treat this lethal transformation.

The High Mobility Group A1 (HMGA1) protein, a non-histone architectural chromatin factor integral to embryogenesis yet typically quiescent in adult tissues, is increasingly recognized for its aberrant re-expression across a spectrum of aggressive solid tumors and high-grade leukemias^16–19^. In such neoplastic settings, HMGA1 functions as a potent chromatin remodeler, orchestrating the activation of critical oncogenic transcriptional programs, including those governed by E2F, MYC, and G2/M cell cycle regulators, with its expression frequently correlating with adverse clinical outcomes^20,21^. Seminal studies in *Jak2*-mutant murine models have implicated HMGA1 as an ‘epigenetic throttle’ in MPN pathogenesis, wherein its genetic attenuation mitigated splenomegaly and myelofibrosis, enhanced ruxolitinib sensitivity, and extended survival^21,22^. Notwithstanding these foundational observations, a comprehensive understanding of HMGA1’s expression dynamics throughout the continuum of MPN progression to sAML, its definitive utility as an early, clinically deployable biomarker, its precise prognostic import, and its exploitability as a therapeutic target in advanced MPN, particularly sAML, remains incompletely elucidated. Building on these antecedent findings, we conducted a comprehensive clinico-genomic investigation, integrating analyses of 240 patient biopsies, single-cell multi-omic datasets, and functional studies employing gain- and loss-of-function leukemia models. This multi-faceted approach was designed to systematically elucidate: (i) The dynamics and quantification of HMGA1 expression, specifically its progressive and clonal upregulation along the MPN-to-sAML continuum in patient-derived materials; (ii) The validation of HMGA1 as a robust, histologically-detectable biomarker with predictive capacity for impending leukemic transformation superior to conventional indices, and its independent prognostic significance for overall survival and treatment outcomes; (iii) The direct functional role of HMGA1 in promoting the aggressive sAML phenotype and mediating resistance to JAK2 inhibitors, and the potential for the next generation JAK2/FLT3 inhibitor pacritinib to counteract such HMGA1-driven oncogenicity. Through the synergistic integration of mechanistic discovery and large-scale patient data interrogation, our overarching objective was to translate an advanced understanding of HMGA1’s pathobiological role into clinically actionable insights that refine risk assessment, facilitate earlier diagnosis, and inform the rational deployment of next-generation therapeutic strategies for high-risk MPN.

## 2 Methods

### 2.1 Patient cohort and Ethical Approval

A total of 240 individuals, including 162 patients with confirmed MPN (essential thrombocythemia, ET, n = 57; polycythemia vera, PV, n = 33; myelofibrosis, MF, n = 51) or MPN-derived sAML (n=21), and 78 patients with suspected MPN ultimately diagnosed with non-MPN hematological conditions (controls), were enrolled at Huashan Hospital, Fudan University; Tongji Hospital, Tongji University; and the First Affiliated Hospital of Naval Medical University (June 2012 - May 2024). Diagnoses were established according to the 5th edition of the WHO classification of hematolymphoid tumors^23^, with central pathology review. The study adhered to the Declaration of Helsinki and received approval from the Institutional Review Boards of Huashan Hospital (No. 2022-1082), Tongji Hospital (No. K-2022-004) and the First Affiliated Hospital of Naval Medical University (No. CHEC2022-076). Written informed consent was obtained from all participants. Convenience sampling was employed for the rare sAML cases. Key inclusion criteria were age 18-90 years with confirmed MPN/sAML and adequate organ function. Exclusion criteria included prior non-hematologic malignancies or confounding comorbidities. Full clinicopathological metadata are provided in Supplementary Table S1, and detailed sample processing is described in the Supplementary Methods.

### 2.2 Cell lines and Culture

Human *JAK2* ^V617F^ mutant (HEL92.1.7, UKE-1), and murine IL-3-dependent (Ba/F3, 32D-cl3) cell lines, including those transduced with *Jak2^WT^*or *Jak2^V617F^* were maintained in RPMI 1640 medium supplemented with 10% fetal bovine serum and 1% penicillin/streptomycin. Murine lines also received 1 ng/mL mouse IL-3. Details of lentiviral transduction for HMGA1/Jak2 overexpression or shRNA-mediated knockdown are in Supplementary Methods

### 2.3 *In*-*vitro* functional assays

Cell viability (CCK8), apoptosis (Annexin V/7-AAD), cell cycle (propidium iodide staining), cytokine-stimulated differentiation (G-CSF treatment of 32D-cl3 cells, CD11b expression), and colony formation assays were performed as described in detail in the Supplementary Methods. Each experiment included at least three biological replicates.

### 2.4 RNA-sequencing (RNA-seq)

Total RNA was extracted (TRIzol), and library preparation (VAHTS Universal V6 RNA-seq Library Prep Kit, Vazyme) was followed by sequencing on an Illumina NovaSeq 6000 (150 bp paired-ends). Reads were processed using fastp, mapped with HISAT2, and differential gene expression was analyzed using DESeq2 (*FDR* < 0.05). Gene Set Enrichment Analysis (GSEA) utilized Hallmark and KEGG gene sets. Detailed bioinformatic pipelines are in Supplementary Methods.

### 2.5 Assay for Transposase-Accessible Chromatin using sequencing (ATAC-Seq)

ATAC-seq was performed on HEL cells transduced with non-targeting control (NC) or HMGA1 shRNA (sh1). Nuclei were isolated, subjected to Tn5 transposition, and DNA was purified. Libraries were prepared (TruePrep DNA Library Prep Kit V2, Vazyme) and sequenced on an Illumina NovaSeq 6000. Raw reads were processed (FastQC, Cutadpat), aligned (Bowtie2), and peaks were called (MACS2). Differential accessibility was determined using DESeq (absolute log2 fold change > 0.5, *FDR* < 0.05). Detaileds are in Supplementary Methods.

### 2.6 Cleavage Under Targets and Tagmentation (CUT&Tag)

CUT&Tag for HMGA1 was performed on HEL and UKE-1 cells transduced with non-targeting control (NC) or HMGA1 shRNA (sh1), using the Hyperactive In-Situ ChIP Library Prep Kit (Vazyme). Briefly, cells were bound to ConA beads, permeabilized, and incubated sequentially with primary HMGA1 antibody, secondary antibody, and pA-Tn5 Transposase. DNA fragments were amplified and sequenced (Illumina NovaSeq 6000). Data analysis involved read processing (fastp), alignment (Bowtie2), peak calling (SEACR), and differential peak analysis (Manorm). Detailed procedures are in Supplementary Methods.

### 2.7 *In vivo* mouse models

Jak2-Flox-V617F/Vav1-Cre-Tg mice (Shanghai Model Organisms Center, Inc.) were used to investigate the impact of *Jak2*^V617F^-driven MPN progression on the aberrant pathological expression of Hmga1. To access the functional consequences of HMGA1 overexpression in MPN-sAML pathogenesis, and to evaluate therapeutic interventions, distinct cohorts of 6-8 week-old NSG mice (Shanghai Model Organisms Center, Inc.) were intravenously injected with luciferase-epxressing HEL cells (transduced with control vector or HMGA1-overexpression constructs). One set of these xenografted mice was used to determine the impact of HMGA1 overexpression on leukemic engraftment, disease progression, and survival. Another set, bearing established HMGA1-overexpressing tumors, was treated with pacritinib (100 mg/kg, BID, orally) or vehicle to assess therapeutic efficacy. Tumor burden was monitored by bioluminescence imaging. Key endpoints included survival, peripheral blood counts, and histopathological analysis of bone marrow and spleen. All animal procedures complied with IACUC guidelines. Detailed experimental designs are provided in the Supplementary Methods.

### 2.8 Omics and Bioinformatics Analysis of Public Datasets

To complement our experimental data and provide a broader context for HMGA1’s multifaceted role in MPN progression to sAML, we conducted extensive bioinformatic analyses of publicly available datasets. These analyses were designed to:

1. Define HMGA1 expression dynamics at single-cell resolution and its association with malignant clones and genetic features during MPN progression to sAML (utilizing single-cell multi-omics datasets: GSE185381, GSE226340, GSE221946).
2. Validate HMGA1 expression trends in larger patient cohorts and relevant murine models, and explore associated molecular pathways (using bulk RNA-sequencing datasets: GSE189979, GSE180851, GSE210253, GSE189570).
3. Investigate the impact of HMGA1 on chromatin accessibility and downstream transcriptional programs, linking its expression to epigenetic and gene regulatory changes (integrating ATAC-seq and RNA-seq data: GSE189570, GSE221946).
4. Assess the clinical and prognostic significance of HMGA1, including its correlation with patient survival and response to therapies in sAML (leveraging the OHSU BeatAML clinical and transcriptomic cohort).
5. Elucidate the differential molecular effects of various JAK inhibitors on HMGA1-associated pathways and resistance mechanisms (analyzing drug-response transcriptomic datasets: GSE229712, GSE190517).

The key bioinformatic tools and specific pipelines applied to these datasets are summarized in Table 1 and detailed in the Supplementary Methods.

**Table 1:** Summary of Publicly Available Datasets and Bioinformatic Analysis Pipelines.

| Pipelines. |  |  |
| --- | --- | --- |
| Dataset | Platform | Key pipeline |
| **Single-cell multi-omics**<br>(GSE185381, GSE226340, GSE221946) | CITE-seq /<br>TARGET-seq/<br>Tapestri single-cell-seq | Seurat v5; Monocle3 v1.2.9; SingCellaR v2.0; Wilcoxon-rank or Kruskal–Wallis with BH; GSEA v4.3 |
| **Bulk RNA-seq** (GSE189979, GSE180851, GSE210253, GSE189570) | Illumina | DESeq2 v1.30.1; paired t-test where applicable. |
| **Drug-response data & clinical outcome data**<br>(GSE229712, GSE190517, OHSU BeatAML) | Illumina/<br>Agilent | DESeq2 v1.30.1; log-rank test, Cox regression; HR-ranked GSEA v4.3. |

### 2.9 Statistical Analysis

Two-group comparisons utilized two-tailed Student’s *t*-test (parametric) or Wilcoxon rank-sum test (non-parametric). Three or more groups were compared using one-way or two-way ANOVA with Tukey’s post-hoc test, or Kruskal-Wallis test with Dunn’s correction. Omics data *P*-values were adjusted for multiple comparisons using the Benjamini-Hochberg method (*FDR* < 0.05). Survival curves were analyzed using the log-rank test, and prognostic factors were assessed by Cox proportional-hazards modeling. IC₅₀ values were derived from four-parameter logistic regression. A *P*-value < 0.05 was considered statistically significant. Analyses were performed using GraphPad Prism v10.0 or R v4.2 (specific package versions: DESeq2 v1.30.1, Seurat v5.0, Monocle3 v1.2.9, SingCellaR v2.0, GSEA v4.3).

### 2.10 Data availability and Code Availability

The in-house generated ATAC-Seq, CUT&Tag and bulk RNA-sequencing data for HEL and UKE-1 with and without HMGA1 knockdown will be deposited in a public repository upon publication. Publicly available datasets used are detailed in Section 2.8 and Supplementary Methods. R scripts used for analyses are available from the corresponding author upon reasonable request.

## 3 Results

### 3.1 HMGA1 is progressively up-regulated during MPN evolution to sAML

To access the breadth and timing of HMGA1 activation, we first analysed single-cell RNA sequencing (scRNA-seq) data from bone marrow mononuclear cells (BMMCs) of three patinets who evlolved from MPN to sAML (17,976 cells) alongside 10 healthy donors (50,244 cells)^24^. *HMGA1* transcripts were markedly enriched in malignant cells (13,763 cells, *FDR* < 0.0001), whereas micro-environmental populations (4,213 cells) displayed baseline expression levels as the healthy donors (Fig. 1a i-iii and Supplementary Fig. 1a). The fraction of *HMGA1*-positive cells increased from a median 19.2% in healthy donors to 46.6% in sAML, and closely tracked the proportion of copy-number-defined malignant clones (Fig. 1a iv; Supplementary Fig. 1b), underscoring disease advanced clone-restricted activation. Validation in two bulk cohorts confirmed a stepwise rise in *HMGA1*: peripheral blood mononuclear cells (PBMCs) and CD34^+^ stem/progenitor cells rose 1.57/2.43-fold from chronic phase (ET/PV) to MF and 1.88/1.58-fold from MF to sAML (both *P* < 0.05; Fig. 1b) and an independent dataset^25^ reproduced this trend (Supplementary Fig. 1c; *FDR* = 0.0378).

**Fig. 1.**
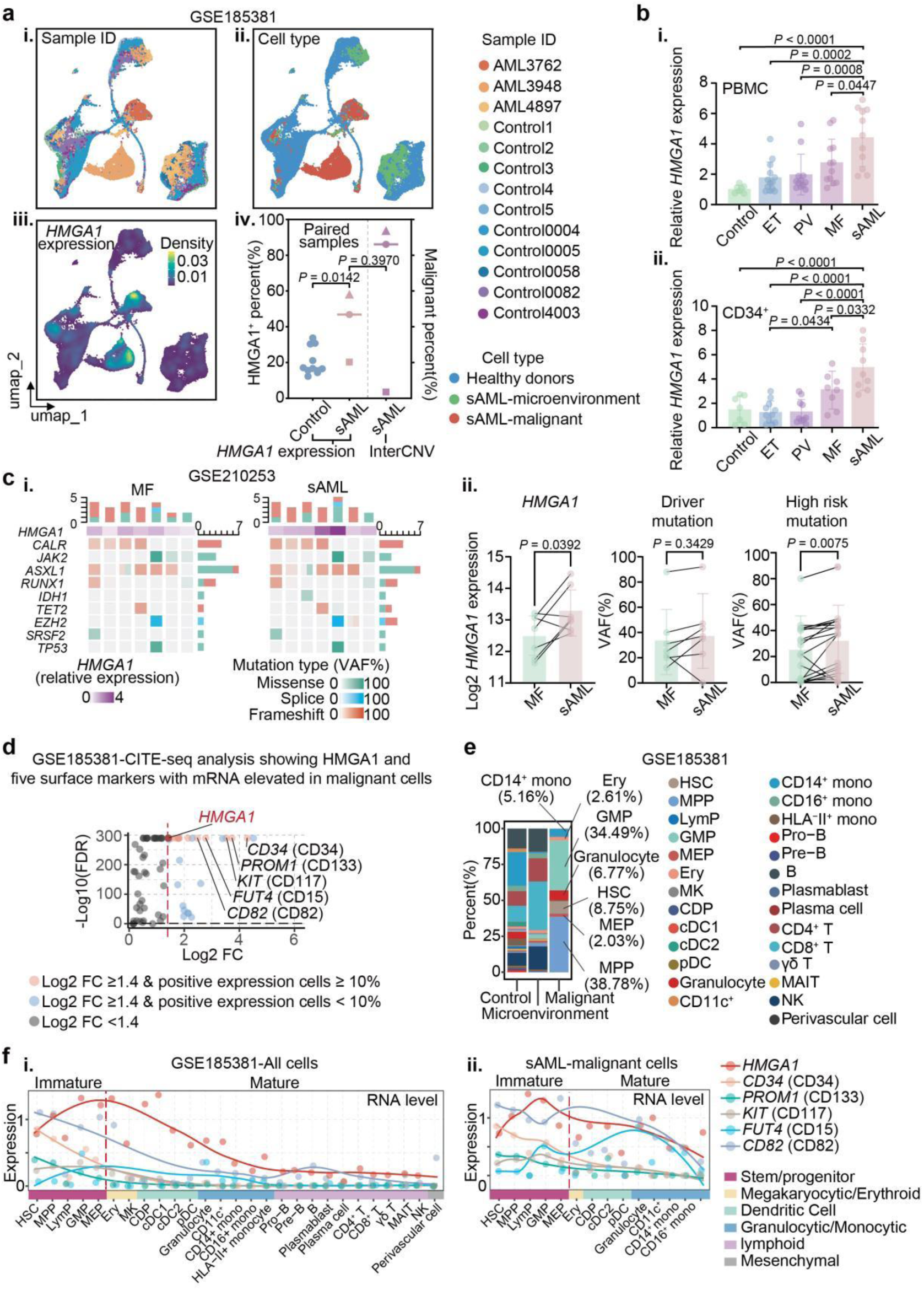
Transcriptomic profiling reveals progressive *HMGA1* upregulation during MPN-sAML evolution. (a) UMAP projections of bone marrow mononuclear cells (BMMCs) from healthy donors (n = 10) and MPN patients progressing to sAML (n = 3) (GSE185381), colored by (i) sample origin, (ii) annotated cell type, and (iii) *HMGA1* transcript expression level. (iv) Correlation plot showing the fraction of *HMGA1*-positive cells versus the fraction of anueploid-define malignant cells in paried MPN-sAML samples. Two-sample *t*-test. (b) Quantitative RT-PCR analysis of *HMGA1* mRNA expression in (i) peripheral blood mononuclear cells (PBMCs; n = 7, 13, 12, 12, 11 for healthy, ET, PV, MF, sAML respectively) and (ii) CD34^+^ cells (n = 7, 11, 10, 8, 9 for healthy, ET, PV, MF, sAML respectively). Data are presented as mean ± standard deviation (SD). One-way ANOVA with Tukey’s post-hoc test. (c) Oncoprint depicting HMGA1 expression and mutational landscape in paired MF and subsequent sAML samples (n = 7 pairs) (GSE210253). (i) Heatmap illustrates relative *HMGA1* expression (color intensity) and mutation burden. (ii) Statistical comparison of *HMGA1* expression, driver mutation load, and high-risk mutation load between MF and sAML phase. Paired two-sample *t*-test. (d) Identifying of five surface markers (CD34, PROM1, KIT, FUT4, CD82) via CITE-seq, whose corresponding mRNA transcripts are significantly elevated in malignant cell populations from sAML patients (GSE185381). (e) Stacked bar chart illustrating the proportions of distinct hematopoietic cell subsets identified by unsupervised clustering of BMMCs from healthy controls and sAML patients (GSE185381). (f) Expression trajectory plot depicting the distribution of transcript expression for *HMGA1*, *CD34*, *PROM1* (CD133), *KIT* (CD117), *FUT4* (CD15), and *CD82* across annotated cell subsets, ordered by developmental hierarchy, for (i) all cells and (ii) malignant cells only.

Paired MF to sAML samples (n = 11)^26^ revealed that escalation of HMGA1 coincided with rising variant allele fractions (VAFs) of high-risk mutations (HRMs), while driver-mutation VAFs remained static (*P* < 0.05; Fig. 1c). To pinpoint which lesions most strongly influence HMGA1 expression, we next leveraged single cell TARGET-seq profiles of 14 *TP53*-mutant sAML cases (9,191 cells). Pseudotime analysis positioned *HMGA1* induction along the trajectory toward proliferative, blast-like state enriched for *TP53* multi-hits, and revealed two HMGA1-high branches with distinct fates (leukemic stem cell-like versus erythroid) (Extended Data Fig. 1a).

Mutation-stratified comparison showed the highest *HMGA1* levels in *TP53*-multi-hit (median normalized counts = 1.39, *FDR* < 0.05) and *JAK2*^V617F^ + *TP53* double-mutant subclones (median normalized counts = 1.61, *FDR* < 0.01), whereas *CALR*-mutant cells displayed much lower expression (median normalized counts = 0, *FDR* < 0.05) (Extended Data Fig. 1b-d). Clones carrying complex karyotypes exhibited the greatest *HMGA1* abundance (Extended Data Fig. 1e, f; *FDR* < 0.0001). These findings implicate HMGA1-mediated expansion of high-risk subclones as a pivotal mechanism driving leukemic evolution through clonal selection.

Because both *JAK2* and *TP53* lesions are prevalent in advanced MPN, we turned to a conditional *Jak2^V617F^*/*Trp53*^R172H^ mouse model^27^. Compared to *Jak2^V617F^* mouse model, *Hmga1* transcripts increased 1.73/1.81-fold in megakaryocyte-erythroid progenitors (MEP) and 1.47/1.74-fold in Lin−Sca-1+c-Kit+ (LSK) stem cells from *Jak2^V617F^*/*Trp53*^R172H/-^ or *Jak2^V617F^*/*Trp53*^R172H/+^ mouse model, whereas complete *Trp53* knockout surprisingly abrogated *Hmga1* upregulation (Extended Data Fig. 1g i-ii; all *FDR* < 0.05). In an advanced MPN-sAML mouse model, MEP cells harboring *Trp53^R172H^* demonstrated even higher *Hmga1* expression (Extended Data Fig. 1h; *FDR* = 0.0060), substantiating that dominant-negative *TP53*, rather than null alleles, synergieses with JAK2 signalling to activate *HMGA1* transcription.

Concordant patterns were noted at the bulk-sample level: PBMCs or CD34^+^ cells bearing *JAK2^V617F^* or obtained from patients who later progressed to sAML consistently exhibited elevated *HMGA1* transcripts than their mutation-negative or non-progressor counterparts (Supplementary Fig. 1d; sAML with *JAK2^V617F^* vs others, *P* < 0.05). *HMGA1* was likewise elevated in sAML cases carrying *TP53* mutations (Supplementary Fig. 1e; PBMC: *P* = 0.0027; CD34^+^: *P* = 0.1864), reinforcing the mutation-stratified single-cell findings.

To examine whether HMGA1 dysregulation extends into mature malignant derivatives, we applied CITE-seq analysis [PMID: 36581735] on BMMCs from 3 sAML alongside 10 healthy donors, simultaneously capturing mRNA and 279 surface proteins. Five surface markers—CD34, PROM1 (CD133), KIT (CD117), FUT4 (CD15) and CD82—showed a transcriptional correlation with malignant identity (Fig. 1d). Unsupervised clustering partitioned malignant cells into multipotent progenitors (MPP; 38.78%), granulocyte– monocyte progenitors (GMP; 34.49%), hematopoietic stem cells (HSC; 8.75%), granulocytes (6.77%), CD14+ monocytes (5.16%), erythrocytes (2.61%), and megakaryocyte-erythroid progenitors (MEP; 2.03%)—with their proportions varying across sAML samples (Fig. 1e and Extended Data Fig. 1i).

Across all cells, the transcription of *HMGA1*, *CD34*, *PROM1*, *KIT*, *FUT4*, and *CD82* gradually declined with increasing maturation state (Fig. 1f i). Focusing on malignant cells alone, however, *HMGA1*, *FUT4*, and *CD82* maintained comparable transcript levels in mature granulocytes and CD14⁺ monocytes versus primitive progenitors (Fig. 1f ii), whereas *CD34* and *KIT* plummeted. However, protein-level validation revealed that FUT4 and CD82 surface abundance decreased in malignant versus non-malignant cells (Supplementary Fig. 1f, g).

Strikingly, HMGA1 transcript remained uniformly high across all leukemic compartments, whereas clinically relevant leukemia markers CD34 and CD117 were confined to primitive subsets (MPP, HSC) and missed in differentiated malignant cells (Extended Data Fig. 1i). Collectively, these data establish *HMGA1* up-regulation as an early and convergent event during MPN progression-most pronounced in *JAK2^v617f^* /*TP53* co-mutant clones and persisting across mature malignant derivates and lay the groundwork for its evaluation as a biomarker of impending leukemic transformation.

### 3.2 HMGA1 as a Diagnostic Biomarker for Leukemic Transformation in MPNs

Having shown that HMGA1 is transcriptionally activated early and across phenotypically diverse malignant cells, we next evaluated its diagnostic utility in patient material. Flow cytometric analysis of 7 diagnostic bone-marrow aspirates revealed a median 86.29% HMGA1^+^ nuclei in blasts, significantly higher than in mature lineages (range 0.79% to 18.56%, *P* < 0.0001; Fig. 2a). Peripheral blood samples showed the same pattern, with sAML cases displaying markedly elevated HMGA1 relative to chronic-phase MPN and healthy donors (*P* < 0.0001; Fig. 2b).

**Fig. 2.**
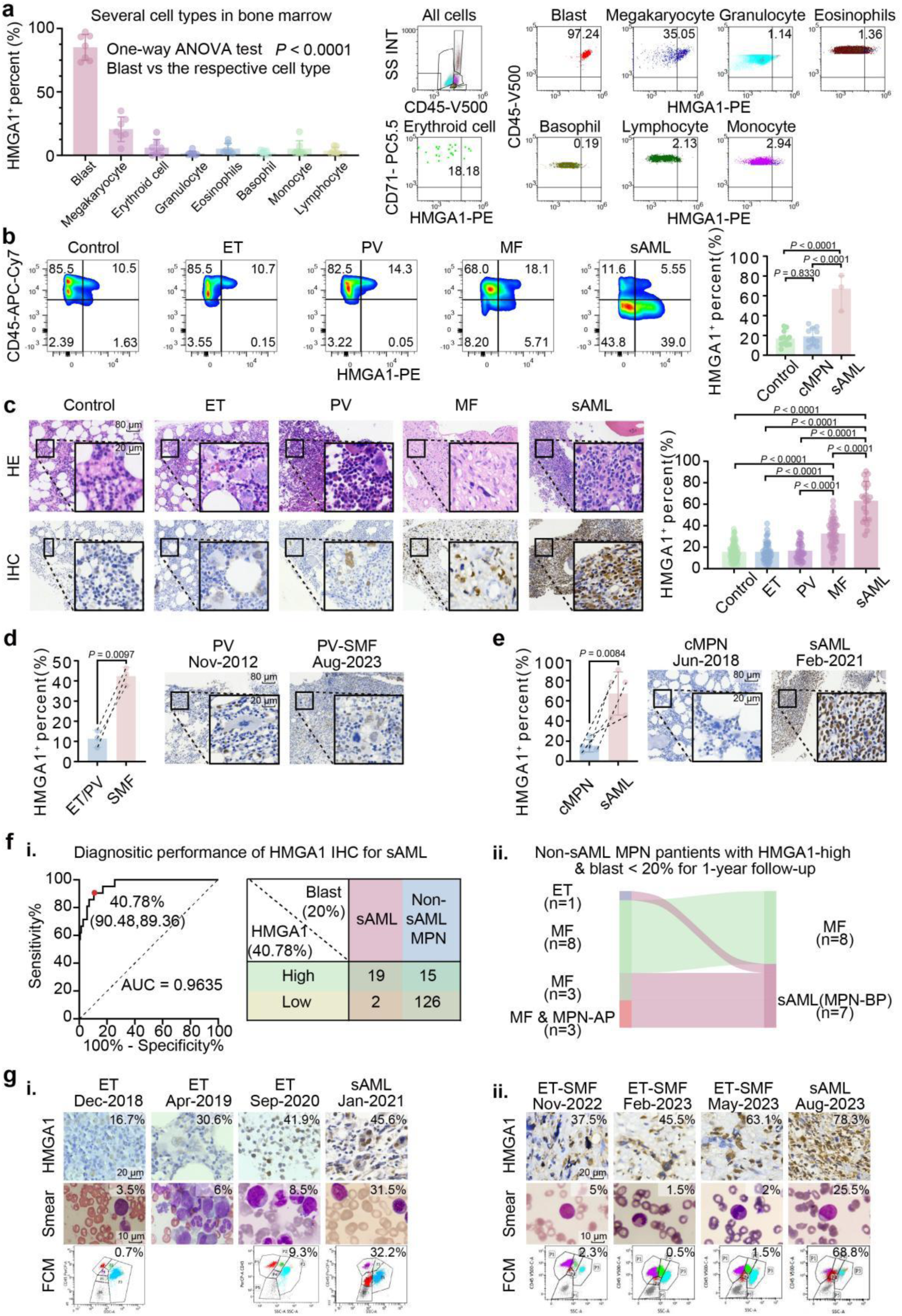
HMGA1 protein as a robust diagnostic biomarker for leukemic transformation in MPN. (a) Flow cytometric quantification of nuclear HMGA1 protein in CD34^+^ blast cells versus matrue hematopoietic lineages from representative MPN patient bone marrow aspirates (n = 7). Data presented as percentage of HMGA1-positive cells. One-way ANOVA with Tukey’s post-hoc test. (b) HMGA1 positive percentage in peripheral blood leukocytes from healthy donors (n = 12), chronic-phase MPN patients (n = 11), and sAML patients (n = 3). One-way ANOVA with Tukey’s post-hoc test. (c) HMGA1 immunohistochemistry (IHC) scores in bone marrow biopsies from patients with ET (n = 57), PV (n = 33), MF (n = 51), sAML (n = 21), and non-MPN controls (n = 78). One-way ANOVA with Tukey’s post-hoc test. Scale bar: top overview, 80 µm; bottom inset, 20 µm. Insets show representative HMGA1 IHC staining. (d, e) Representative HMGA1 IHC staining in paried bone marrow biopsies from patients at chronic phase MPN versus post-myelofibrosis (post-MF, n = 3) and (e) chronic phase MPN vsrsus sAML transformation (n = 5). Scale bars as in (c). Paired two-sample *t*-test. (f) (i) Receiver operating characteristic (ROC) curve analysis of of HMGA1 IHC score for discriminating sAML (n = 21) from non-sAML MPN (n = 141). Area under the curve (AUC) is indicated. Right panel: contingency table at the optimal HMGA1 cut-off. (ii) Sankey diagram illustrating the 1-year clinical course of 15 <u>n</u>on-blast-phase MPN patients with high baseline HMGA1 expression (> 40.78%) but < 20% bone marrow blasts. (g) Longitudinal analysis of two representative MPN patients who progressed to sAML. Serial bone marrow biopsy images (HMGA1 IHC, scale bar: 10 µm); bone marrow smear cytology (Wright-Giemsa staining, scale bar: 20 µm); and corresponding flow cytometry dot plots (CD45 vs. side scatter) at indicated time points, demonstrating HMGA1 upregulation and moprpholoigcal changes preceding overt blast crisis.

A multi-centre tissue microarray comprising 240 bone-marrow biopsies—78 from pateints whose marrow samples were ultimately non-MPN (served as disease controls) and 162 pathologically confirmed MPN comprising ET (n = 57), PV (n = 33), MF (n = 51) and sAML (n = 21)—confirmed a step-wise increase in nuclear HMGA1 staining (ET/PV 16.03 % ± 8.43, MF 32.58 % ± 12.88, sAML 63.21 % ± 18.04; *P* < 0.0001; Fig. 2c). Sporadic megakaryocytic HMGA1 up-regulation was noted in 7/90 ET/PV cases, hinting at early focal activation. Paired longitudinal biopsies demonstrated an average 3.74/4.37-fold rise in HMGA1 on progression to post-MF (sMF) or sAML (*P* < 0.01; Fig. 2d, e).

Across the cohort, HMGA1 levels were higher in patients with splenomegaly (*P* < 0.0001) and in those with a diagnostic latency ≥ 10 months (*P* = 0.0001) (Extended Data Fig. 2a, b). HMGA1 tracked positively with reticulin grade (MF-0 vs MF-2/3; *P* < 0.01) and inversely with hemoglobin, hematocrit and platelet counts (all *P* < 0.0001) but not total white-cell count (Extended Data Fig. 2c, d). Expression was independent of sex, age or IPSS-risk category (Supplementary Fig. 2a-e), yet adverse ELN sAML carried the highest HMGA1 (Extended Data Fig. 2e).

Genotyping showed that *JAK2*^V617F^ and *TP53* mutations correlated with elevated HMGA1 in both chronic-phase MPN and sAML (Supplementary Fig. 2f, g), whereas other driver or high-risk lesions did not. Interestingly, apart from the *TP53* mutations, other driver and high-risk mutations were not significantly enriched in the sAML cohort, suggesting that leukemia transformation cannot be reliably predicted based on these mutations alone (Supplementary Table 4). In a *Jak2*^V617F^ knock-in mouse, *Hmga1* transcripts and megakaryocyte expansion paralleled the human data (*P* = 0.0041; Supplementary Fig. 2h-j).

Using sAML (≥20% blasts) as outcome, HMGA1 IHC achieved AUC = 0.96 (95% CI 0.94-0.99), sensitivity 90.48% and specificity 89.36% at a 40.78% cut-off (Fig. 2f i). Among 15 chronic-phase MPN patients with high HMGA1 expression but <20% blasts, 7 (ET n=1; MF n=3; MF & MPN-AP n=3) converted to sAML within 12 month (median lead time 6 months, IQR 4-9 months; Fig. 2f ii).

Serial smears from two representative converters revealed degenerative granulocytic morphology (loss of neutral particles, particle thickening, vacuolization; pathological hallmarks correlating with early leukemic transformation) that coincided with HMGA1-high expansion and preceded overt blast escalation (Fig. 2g), indicating that HMGA1 marks molecular progression ahead of conventional criteria.

*HMGA1* was highly expressed in both MPN-derived sAML lines—HEL (FAB-M6, CD34^+^CD117^-^) and UKE-1 (FAB-M7, CD34^+^CD117^+^)—underscoring its subtype-agnostic nature (Supplementary Fig. 2k,l). Multiplex immunofluorescence (mIF) of patient biopsies showed that HMGA1 localized to megakaryocytes in chronic MPN but shifted to blasts in sAML, while CD34/CD117 were variably expressed and largely confined to primitive niches (Extended Data Fig. 2f-j and Supplementary Fig. 2m). Co-staining in an advanced sAML sample demonstrated near-complete overlap of HMGA1 with both CD34 and CD117 in blasts (Supplementary Fig. 2n, o).

Longitudinal tracking in a patient evolving from ET to sMF, and ultimately to sAML revealed that HMGA1 rose first from ET to sMF; CD34 increased mainly within neovascular regions, and CD117 spiked only at frank transformation (Extended Data Fig. 2k). Thus, HMGA1 provides broader and earlier coverage of malignant clones than CD34 or CD117, rationalizing its superior ROC performance.

Together, these findings nominate HMGA1 IHC staining in bone-marrow biopsies, particularly when interpreted alongside *JAK2*^V617F^ and *TP53* status, as an actionable tool for early detection of leukemic progression and could aid in early identification of patients at risk for sAML transformation.

### 3.3 HMGA1 Promotes Clonogenic Growth, Blocks Differentiation and Accelerates Leukemia progression by Directly Modulating Core cell Cycle Machinery

While previous studies have implicated HMGA1’s association with MPN cell proliferation^21^, its specific role in driving sAML progression, particularly the *in vivo* consequences of its overexpression and its precise influnece on hematopoietic differentiation, required further elucidation. We therefore sought to comprehensively define the functional consequences of elevated HMGA1 in sAML and determine whether it directly contributes to malignant behavior. To this end, we engineered stable *HMGA1* knock-down (sh1, sh2; > 80% depletion) and over-expression (OE; >2-fold increase) in four different MPN/sAML cell lines—human HEL and UKE-1, and murine Ba/F3 and 32D-cl3 (Supplementary Fig 3a-c). Phenotypes were consistent across both shRNAs, confirming on-target effects.

HMGA1 overexpression robustly enhanced colony-forming unites (CFU) 1.36-fold in HEL and 1.31_fold in UKE-1, whereas knock-down reduced CFU by 46/61% (sh1) and 22/41% (sh2) (all *P* < 0.01; Fig 3a). Growth-curve assays confirmed these CFU findings, with OE cultures expanding more rapidly than controls and KD cultures showing reduced proliferation (Extended Data Fig. 3a). Importantly, Hmga1 overexpression did not render Ba/F3 cells cytokine-independent, indicating that external growth cues remain obligatory (Extended Data Fig. 3a). In 32D-cl3 cells, Hmga1 overexpression suppressed G-CSF-induced granulocytic maturation, blocking CD11b up-regulation and morphological differentiation (all *P* <0.0001; Extended Data Fig. 3b). Together, these data show that HMGA1 simultaneously boosts proliferation and maintains an undifferentiated state without bypassing cytokine requirements.

**Fig. 3.**
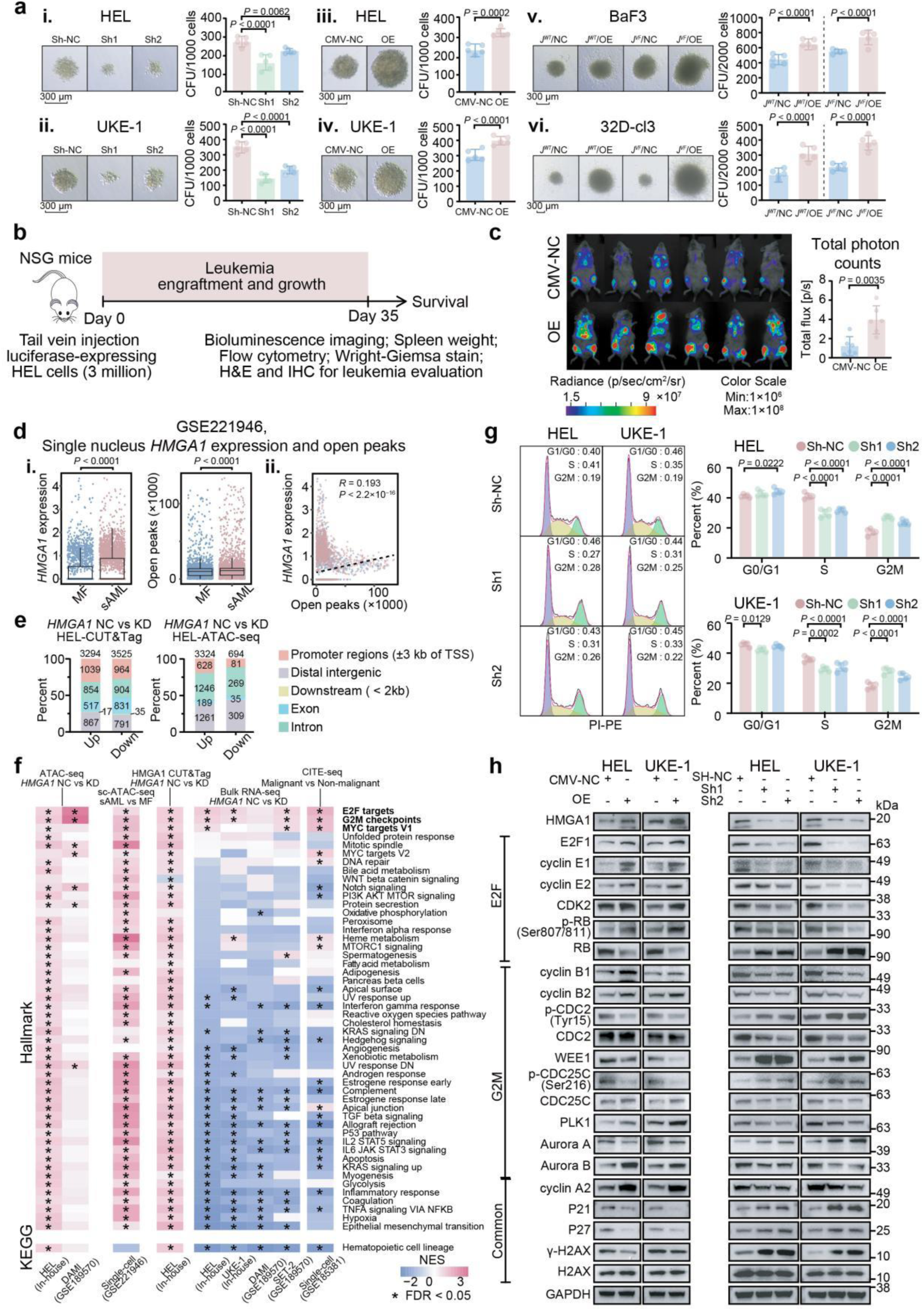
HMGA1 promotes pro-leukemic phenotypes and directly regulates cell cycle machinery via chromatin modulation. (a) Colony-forming units (CFU) assays in human (HEL, UKE-1) and murine (Ba/F3-*JAK2*^V617F^, 32D-*JAK2*^V617F^) cell lines following HMGA1 knockdown (shHMGA1) or overexpression (OE-HMGA1) relative to respective controls (NC). Data are mean ± SD. (n=5 per group). Representative colony images are shown (scale bar 300 μm).Two-sample *t*-test and one-way ANOVA with Tukey’s post-hoc test. (b) Schematic of the HEL cell xenograft experiment in NSG mice, comparing HMGA1-OE cells to vector control cells. (c) Representative bioluminescence images (left panel) and quantification of total photon flux (right panel) from NSG mice at day 35 post-engraftment with luciferase-tagged HEL cells transduced with vector control (CMV-NC, n=6) or HMGA1 overexpression constructs (OE, n=6). Data are mean ± SD. Two-sample *t*-test. (d) Correlation between *HMGA1* mRNA expression (snRNA-seq) and the number of open chromatin peaks (scATAC-seq) in CD34^+^ cells from a patient during primary MF and after sAML transformation (GSE221946). Pearson correlation coefficient (*R*) and *P*-value from Wilcoxon ranksum test with BH correction. (e) Genomic distribution of HMGA1 binding sites lost upon HMGA1 knockdown in HEL cells, as determined by CUT&Tag (i), and chromatin regions with significantly reduced accessibility upon HMGA1 knockdown, as determined by ATAC-seq (ii). Stacked bar chats show the percentage of peaks located in promoter (±3kb of TSS), intronic, intergenic, and other genomic regions. (f) GSEA enrichment plots for E2F targets and G2M checkpoint Hallmark gene sets. Comparisons shown in figure. Normalized Enrichment Score (NES) and False Discovery Rate (FDR) q-value are indicated. (g) Cell cycle distribution (Propidium Iodide staining followed by flow cytometry) of HEL and UKE-1 cells with HMGA1 knockdown (sh1, sh2) or non-targeting control (NC). Data are mean ± SD. (n = 5 per group). One-way ANOVA with Tukey’s post-hoc test. (h) Western blots analysis of key E2F pathway, G2M checkpoint, and common cell cycle regulatory protein levels in HEL and UKE-1 cells following overexpression (OE vs. CMV-NC vector control; left panels) or HMGA1 knockdown (shHMGA1 vs. shNC control; right panels). GAPDH served as loading controls. Blots are representative of at least three independent experiments.

To determine the *in vivo* impact of HMGA1 overexpression, we transplanted luciferase-labelled HEL cells (OE or vector) intravenously into NSG mice (schema, Fig. 3b). Longitudinal bioluminescence imaging revealed a steeper growth curve in HMGA1-OE recipients, reaching a 3.27-fold higher total bioluminescent by day 35 compared with controls (Fig. 3c). Peripheral-blood flow cytometry and Wright–Giemsa smears consistently showed elevated circulating blasts in HMGA1-OE mice (Extended Data Fig. 3c, d). OE markedly increased leukemic engraftment in bone marrow and extramedullary sites. Histology of the bone-marrow showed densely packed HMGA1-positive cells (Extended Data Fig. 3e) and diffuse HMGA1-positive infiltration in spleen (Extended Data Fig. 3f). Spleen weight has increased by 43.0% (20.2% vs 28.8% of body weight, *P* = 0.0398; Extended Data Fig. 3g). HMGA1-OE shortened median survival from 45.5 to 36 days (hazard ratio = 4.10, log-rank *P* = 0.0022; Extended Data Fig. 3h). Of note, peripheral blood counts and body weight trajectories did not differ (Supplementary Fig 3d), indicating death was driven by tumor burden rather than systemic toxicity. Together, these data demonstrate that enforced HMGA1 expression enhances leukemic dissemination, promotes splenic and marrow infiltration, and significantly worsens survival *in vivo*.

To elucidate the molecular underpinnings of HMGA1-driven leukemogenesis, we first examed HMGA1’s impact on the chromatin landscape. Analysis of published scATAC-seq data from paired sAML patient samples undergoing transformation^28^ revealed a positive correlation between *HMGA1* upregulation and an increase in open chromatin peaks (*R* = 0.193, *P* < 2.2×10^16^; Fig. 3d), suggesting a role for HMGA1 in in modulating chromatin accessibility during disease progression. Prompted by this, we performed CUT&Tag sequencing for HMGA1 in HEL, alongside ATAC-seq in HEL cells, comparing non-targeting control (NC) with HMGA1 shRNA (sh1) conditions. CUT&Tag analysis identified 20,511 high-confidence HMGA1 binding peaks in control HEL cells, of which 3,294 were significantly diminished upon HMGA1 knockdown (*FDR* < 0.05); a substantial fraction (31.5%) of these lost sites localized to promoter regions (±3 kb of TSS) (Fig. 3e and Extended Data Fig. 3i). Critically, these HMGA1-depeted binding sites were highly enriched at the regulatory elements of core cell cycle genes, including G1/S phase regulators (*E2F1*, *CCNE1*, *CCNE2*, *CDK2*, *RB1*), and G2/M phase drivers (*CCNB1*/*2*, *CDC2*, *WEE1*, *CDC25C*, and *AURKB*) (Supplementary Fig. 3e), indicating direct HMGA1 occupancy crucial for their regulation.

Concordantly, ATAC-seq in HEL cells demonstrated that HMGA1 knockdown resulted in significantly reduced accessibility at 3,324 chromatin regions (Fig. 3e and Extended Data Fig. 3i). A significant overlap was observed between regions losing HMGA1 binding (CUT&Tag) and those exhibiting decreased accessibility (ATAC-seq), particularly at loci enriched for E2F-Cyclin-CDK axis genes (Supplementary Fig. 3e). This direct linkage underscores HMGA1’s role in maintaining a permissive chromatin state at these cell cycle gene promoters. Both lost CUT&Tag peaks and differentially accessible ATAC-seq regions were significantly enriched for genes within E2F and G2/M checkpoint pathways (Fig 3f). Further supporting a model of specific chromatin modulation, only 694 regions gained accessibility upon HMGA1 knockdown, predominantly in distal intergenic areas (Fig. 3e). Motif analysis of regions losing accessibility upon HMGA1 depletion revealed a concomitant loss of AP-1, ETS, and NF-κB transcription factors, whereas regions gaining accessibility were enriched for CTCF/BORIS and IRF motifs (Extended Data Fig. 3j), suggesting HMGA1 may facilitate the binding of pro-proliferative transcription factors by displacing nucleosomes. This regulatory function appeared conserved, as *HMGA1* knockdown in DAMI cells also led to reduced chromatin accessibility at thousands of peaks (4,803 lost), with a notable proportion (∼8.8%) in promoters, and was associated with enrichment of E2F and G2/M pathways (Fig 3f). Furthermore, analysis of scATAC-seq from paired sAML patient samples^28^ corroborated HMGA1’s pivotal role in maintaining chromatin accessibility at E2F and G2/M pathway-associated regions during leukemic transformation (Fig 3f).

Consistent with these profound chromatin alterations, GSEA of RNA-seq data from CITE-seq of malignant versus non-malignant patient cells^24^ and our HMGA1-pertubed cell lines (HEL, UKE-1), and public data (DAMI, and SET-2)^21^ consistently highlighted E2F targets, MYC targets and G2/M checkpoints pathways as the most significantly activated Hallmark gene sets in HMGA1-proficient context (NES > 1.2, *FDR* < 0.01, except DAMI *FDR* < 0.1), whereas KD restored hematopoietic programs (Fig. 3f; Supplementary Fig. 3f).

These transcriptional changes translated directly to cellular phenotypes: PI profiling in HEL cells showed that HMGA1-KD reduced the S-phase fraction (41.2 % to 30.0/31.7 %) and increased G2/M arrest (17.3 % to 27.2/23.6 %) (Fig. 3g). Annexin-V/7-AAD staining demonstrated a two-fold rise in apoptosis after knockdown in HEL (9.12 % vs 15.51/19.29 %, *P* < 0.0001; Extended Data Fig. 3k).

Immunoblotting provided the biochemical underpinning for these effects (Fig. 3h). HMGA1 overexpression orchestrated a classic G1/S-activation signature (increased E2F1, cyclin E1/E2 and CDK2; hyper-phosphorylated RB; decreased CDK inhibitors P21 and P27) and facilitated G2/M propgression (increased PLK1, Aurora B, cyclin B1/2, cyclin A2; suppressed checkpoint kinase WEE1 and inhibitory p-CDK1). Conversely, HMGA1 knockdown reversed these changes and induced γ-H2AX accumulation, indicative of replication stress and G2/M checkpoint enforcement.

Collectively, our gain- and loss- of-function data supported by detailed chromatin analyses, show that HMGA1 orchestrates chromatin opening at core cell cycle loci, thereby accelerating cell-cycle progression and promoting survival pathways that drive clonogenic expansion *in vitro* and aggressive disease *in vivo*.

### 3.4 HMGA1 predicts JAK-Inhibitor Resistance and Prognosis in MPN-sAML

Having established HMGA1 as a mechanistic driver of aggressive disease biology, we next asked whether its expression stratifies clinical outcomes and therapy response in patient cohorts.

In the OHSU BeatAML discovery cohort (n = 31 MPN-sAML), HMGA1 ranked within the top 0.2 % (65/22,837) of genes most strongly associated with inferior overall survival (HR = 3.04, 95 % CI 1.26-7.35; log-rank P = 0.0096; Fig. 4a). Gene-set enrichment on HR-ranked genes highlighted E2F, G2/M and MYC programs (*FDR* < 0.05; Fig. 4b), mirroring the mechanistic data in Section 3.3.

**Fig. 4.**
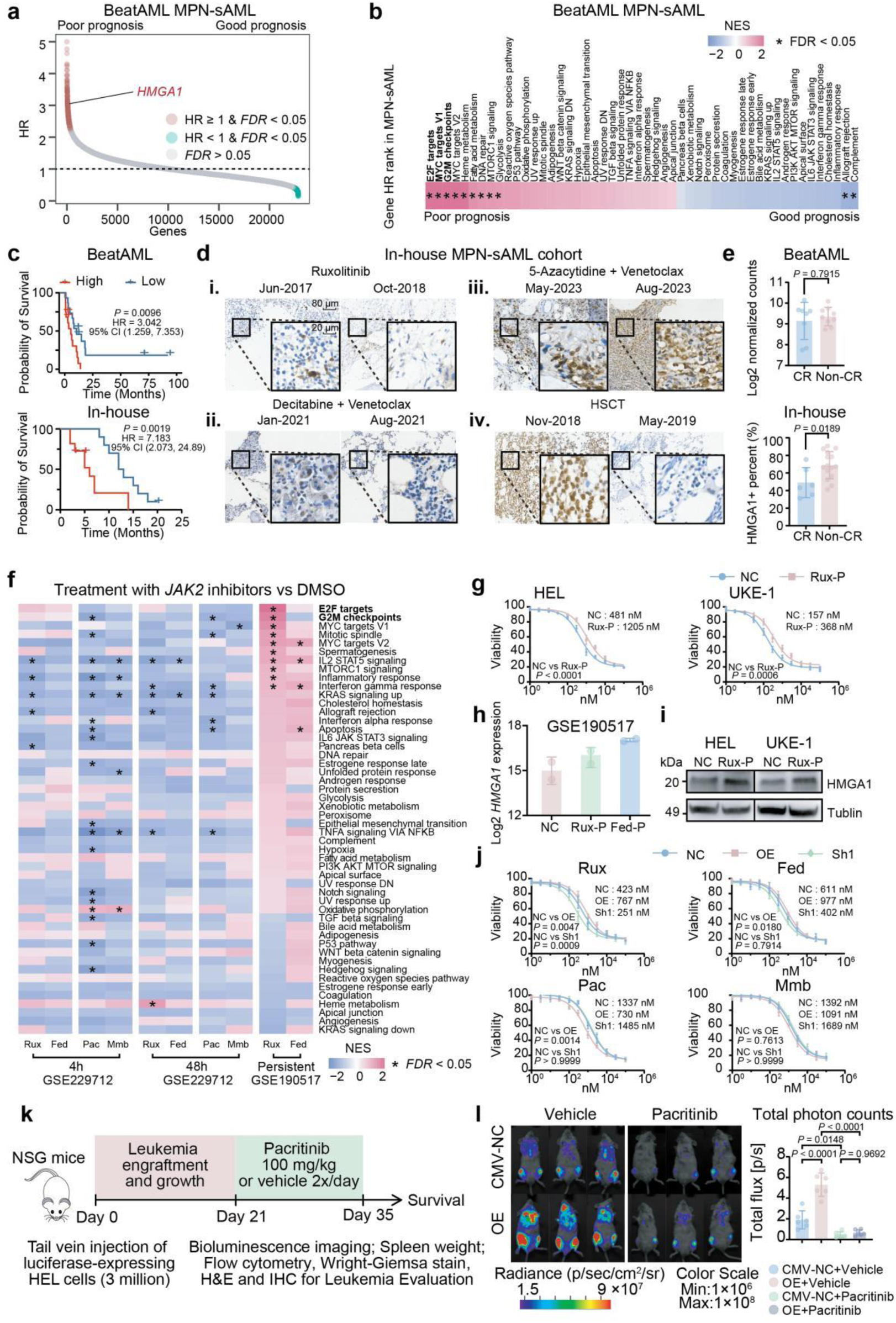
HMGA1 expression dictates clinical prognosis and therapeutic response to JAK inhibitor5s, with pacritinib overcoming HMGA1-mediated resistance. (a) Prognostic significance of *HMGA1* expression in the OHSU BeatAML MPN-sAML cohort (n = 31). Genes are ranked by their hazard ratio (HR) for overall survival (OS). Points are colored based on *FDR* significance: grey (*FDR* > 0.05), blue (*FDR* < 0.05 & HR < 1, good prognosis), red (*FDR* < 0.05 & HR ≥ 1, poor prognosis) (b) Gene Set Enrichment Analysis (GSEA) showing Hallmark pathways enriched among genes associated with poor prognosis (ranked by HR) in the OHSU BeatAML MPN-sAML cohort. Heatmap displays NES for selected pathways. Asterisks (*) indicate *FDR* < 0.05. (c) Kaplan-Meier OS curves for MPN-sAML patients from the OHSU BeatAML cohort (n = 31, top panel) and an independent in-house cohort (n = 21, bottom pnel), stratified by high versus low *HMGA1* expression (HMGA1 expression levels for BeatAML in-house cohort using median cut-off). Log-rank (Mantel-Cox) test. (d) Representative immunohistochemical (IHC) staining for HMGA1 in bone marrow biopsies from in-house MPN-sAML cohort patients, illustrating HMGA1 expression changes with therapy and clinical outcome. (i) HMGA1-low patient achieving complete remission (CR) post-ruxolitinib. (ii) HMGA1-low patient achieving CR post-decitabine + venetoclax. (iii) HMGA1-high patient with progressive disease (PD) despite 5-azacytidine + venetoclax, showing increased HMGA1 at relapse. (iv) HMGA1-high patient achieving durable remission with decreased HMGA1 staining post-allogeneic hematopoietic stem cell transplantation (allo-HCT). Scale bars: 80µm (overview), 20µm (insets). (e) Comparison of HMGA1 expression levels between MPN-sAML patients achieving complete remission (CR, includes CRh, CRi) and those not achieving CR (Non-CR). Top panel: HMGA1 transcript levels (Log2 normalized counts) in the OHSU BeatAML cohort (n=31). Bottom panel: Percentage of HMGA1-positive cells (IHC score) in the in-house cohort (n=21). Data are mean ± SD. Two-sample *t*-test. (f) Heatmap illustrating Hallmark GSEA of differentially expressed genes in HEL cells treated with DMSO (vehicle), ruxolitinib (Rux), fedratinib (Fed), pacritinib (Pac), or momelotinib (Mmb) for 4 hours or 48 hours (GSE229712) and in HEL cells with acquired resistance to ruxolitinib (Rux-Persistent, GSE190517) compared to DMSO control. Color intensity represents NES. * indicate *FDR* < 0.05. (g) Dose-response curves depicting cell viability of parental (NC) control versus ruxolitinib-persistent (Rux-P) HEL (left) and UKE-1 (right) cells treated with indicated concentrations of ruxolitinib for 72 hours. IC_50_ values (mean± SD) are shown. Two-way ANOVA comparing IC_50_ values. (h) *HMGA1* mRNA expression (RNA-seq, normalized counts) in HEL cells: non-targeting control (NC), ruxolitinib-persistent (Rux-P), and fedratinib-persistent (Fed-P). (I) Immunoblot analysis of HMGA1 protein levels in parental (NC) and and ruxolitinib-persistent (Rux-P) HEL and UKE-1 cells. Tublin serves as a loading control. (J) Dose-response curve showing cell viability of HEL cells transduced with control vector (NC), HMGA1 overexpression (OE), or HMGA1 shRNA (Sh1) constructs, treated with indicated concentrations of ruxolitinib (Rux), fedratinib (Fed), pacritinib (Pac), and momelotinib (Mmb) for 72 hours. IC50 values (mean ± SD) are shown. Two-way ANOVA comparing IC50 values between OE/Sh1 and respective NC. (k) Schematic representation of the in vivo pacritinib treatment study in NSG mice engrafted with luciferase-expressing HEL cells (transduced with CMV-NC vector or HMGA1-OE construct). Following leukemia engraftment (Day 0-21), mice received pacritinib (100 mg/kg, BID) or vehicle orally for 14 days (Day 21-35). Endpoint analyses included bioluminescence imaging, spleen weight, flow cytometry, Wright-Giemsa staining, H&E, and IHC, alongside survival monitoring. (l) Representative bioluminescence image (left) and quatification of total tumor bioluminescence (total flux, right) at day 35 in NSG mice engrafted with CMV-NC or HMGA1-OE HEL cells and treated with vehicle or pacritinib (n=6 mice/group). Data are shown in mean ± SD. One-way ANOVA with Tukey’s post-hoc test.

Validation in an independent, in-house sAML cohort (n = 21) confirmed that HMGA1-high patients had a median OS of 5 months versus 12 months in HMGA1-low (HR = 7.18, 95% CI 2.07-24.89; log rank *P* = 0.0019; Fig 4c). Consistenly, multivariable Cox analysis confirmed HMGA1-high as the sole independent predictor of poor overall survival (BeatAML HR = 3.02, 95% CI 1.16-7.88; log-rank *P* = 0.0236; in-house HR = 6.69, 95% CI 1.62-27.62; log-rank *P* = 0.0087), whereas patient age, sex and ELN-2022 risk category were not significant (Extended Data Fig. 4a).

Standard-of-care regimens including ruxolitinib, hypomethylating agents (HMA), cytarabine and hydroxyurea, afforded no survival advantage en bloc (Extended Data Fig. 4b). HMGA1 stratification revealed sharp heterogeneity in treatment response patterns: 4/6 HMGA1-low cases achieved complete remission (CR) with ruxolitinib or decitabine-venetoclax without secondary HMGA1 induction (Fig. 4d i-ii), where as HMGA1-high patients progressed despite 5-azacytidine-venetoclax and even further up-regulated HMGA1 (Fig. 4d iii). One HMGA1-high patient attained durable remission only after allogeneic transplantation, which reduced HMGA1 staining by 70 % (Fig. 4d iv), implying that eradication of the malignant clone can eliminate the high-HMGA1 phenotype.

Across both cohorts, high HMGA1 was linked to lower CR or complete remission with incomplete hematologic recovery (CRi) rates with mpn-sAML-related regimens (BeatAML *P* = 0.7915; in-house *P* = 0.0189) (Fig. 4e; Supplementary Fig. 4a-f). These clinical trends align with *in vitro* data in Section 3.5, where HMGA1-OE lines exhibited primary resistance to current treatments, especially in first-generation type-I JAK2 inhibitors-ruxolitinib.

In short, high HMGA1 warns of poorer prognosis and identifies tumours that are unlikely to respond to current treatments. These observations underscore the need for therapeutic strategies that can bypass or neutralize HMGA1-mediated resistance.

### 3.5 Pacritinib Suppresses HMGA1-Mediated Disease Burden and Progression

Given the limited therapeutic options for HMGA1-high sAML and the absence of direct HMGA1 inhibitors, we next sought alternative strategies to target HMGA1-driven pathways. We compared four JAK2-targeted agents with distinct generational and selectiveity profiles: the first-generation JAK inhibitor ruxolitinib; and three next-generation inhibitors-fedratinib (more JAK2-selective), pacritinib (JAK2/FLT3/IRAK1 inhibitor), and momelotinib (JAK1/JAK2/ACVR1 inhibitor)—for their ability to modulate HMGA1 expression and downstream signaling in HEL cells, using RNA-seq (GSE190517^29^ and GSE299712^30^) and protein analyses after short (4 hr) and prolonged (30 day) exposure.

Brief (4 hr) exposure to any of the inhibitors initially reduced STAT3/5 phosphorylation and broadly down-shifted proliferative programs. However, transcriptomic analysis by GSEA revealed a rapid divergence in their effects on key oncogenic pathways. By 4 hours, ruxolitinib and fedratinib permitted the reactivation of E2F targets and G2/M checkpoint gene sets. In contrast, pacritinib, with its distinct multi-kinase profile including FLT3 and IRAK1 inhibition alongside JAK2, uniquely mainained suppression of these pathways, even though HMGA1 mRNA levels remained unchanged at this early time point. Momelotinib showed an intermediate profile (Fig. 4f).

Prolonged (30-day) exposure to ruxolitinib and fedratinib highlighted the development of adaptive resistance. This was evidenced by a markedly increased drug IC_50_ for ruxolitinib (HEL: 481 nM vs 1205 nM; UKE-1: 157 nM vs 368 nM Fig. 4g) and was accompanied by an upregulation of both HMGA1 mRNA (Fig. 4h) and protein (Fig. 4i) specifically in ruxolitinib-persistent cells. GSEA further confirmed that persistent ruxolitinib or fedratinib treatment led to the sustained re-activation of E2F targets and, for ruxolitinib, also G2/M checkpoint genes (Fig. 4f, “persistent” tracks). This transcriptional reprogramming towards a proliferative state, despite initial inhibition, underscores a key mechanism of acquired resistance to these inhibitors.

At the protein level, pacritinib’s distinct mechanism was further evident. It sustained the inhibition of p-STAT3/p-STAT5 and effectively down-regulated key cell cycle drivers including E2F1, cyclin E/CDK2, and PLK1. Concurrently, pacritinib up-regulated crucial cell cycle checkpoint proteins (RB, WEE1, p-CDC25C, p21, p27) and induced the DNA damage marker γ-H2AX (Extended Data Fig. 4c), indicative of cell cycle arrest and induction of apoptosis. In stark contrast, cultures treated long-term with ruxolitinib exhibited a rebound of proliferative drivers like E2F1 and cyclin B2, with a concomitant reduction in γ-H2AX. This, coupled with the GSEA findings showing that long-term ruxolitinib exposure leads to further activation of E2F and G2/M-related genes, suggests an engagement of adaptive DNA repair mechanisms that promote cell survival and resistance (Supplementary Fig. 4g).

Taken together, these transcriptomic and proteomic data position pacritinib as a durable suppressor of HMGA1-associated cell-cycle circuitry, acting effectively downstream of, or parallel to, HMGA1 transcription, and capable of overcoming the adaptive resistance observed with other JAK2 inhibitors like ruxolitinib and fedratinib.

Dose-response assays in HEL, UKE-1, Ba/F3 and 32D-cl3 lines confirmed that HMGA1- OE increased IC₅₀ values for ruxolitinib and fedratinib by 1.2- to 2-fold, while HMGA1-KD decreased them by 12.6-46.6% (*P* < 0.01). Pacritinib’s IC₅₀ (48∼730 µM) decreased by 18.3-45.4% across HMGA1-OE backgrounds, and momelotinib displayed only modest efficacy, especially in OE clones (Fig. 4j; Supplementary Fig. 4h-j). These findings further underscore that pacritinib retains its efficacy irrespective of HMGA1 expression status, unlike the other inhibitors, which are significantly impacted by HMGA1 levels.

We next evaluated the antileukemic efficacy of pacritinib in the HMGA1-OE NSG xenograft model (treatment schema; Fig. 4k). After confirming engraftment on day 21, mice received oral pacritinib (100mg/kg, BID) or vehicle for 14 days. Pacritinib lowered circulating CD45^+^CD117^+^ blasts from 6.92 % ± 1.82 to 1.40 % ± 0.66 and reduced Wright-Giemsa-counted HEL cells in blood smears by 60% (all *P* < 0.0001; Extended Data Fig. 4d). Drug-treated mice showed >50% fewer HMGA1-positivie cells in bone-marrow and spleen (Extended Data Fig. 4e, f). Correspondingly, spleen weight fell from 0.34 % to 0.22 % of body weight (*P* =0.0023) and bioluminescent tumor flux dropped 8.0-fold (Fig. 4g). Pacritinib doubled median survival from 38 days (vehicle) to 79 days (log-rank *P* = 0.0006) in HMGA1-OE recipients, approaching survival of vector-control (non-HMGA1-overexpressing) given pacritinib (Extended Data Fig. 4h). The drug alleviated cachexia-associated weight loss (*P* = 0.0212; supplementary Fig. 4k) without altering peripheral counts (*P* > 0.05; Supplementary fig. 4l).

These integrated *in vitro* and *in vivo* finding position pacritinib, a clinically approved next-generation JAK2/FLT3/IRAK1 inhibitor as a rational treatment for HMGA1-high MPN-sMAL, particularly after failure of other JAK2 inhibitors. Incorporating HMGA1 status into therapeutic decision-making could help identify patients most likely to benefit from pacritinib and spare others from ineffective first-generation agents.

## 4. Discussion

Our findings substantially reinforce and expand the understanding of HMGA1 as a central regulatory node in MPN-sAML progression. This study delineates HMGA1 not only as a key mechanistic driver of leukemic transformation, particularly in synergy with canonical *JAK2*^V617F^ and *TP53* lesions, but also establishes its significant translational value as a superior, IHC-detectable biomarker for predicting sAML development and stratifying patient prognosis. Critically, we unveil HMGA1 as a therapeutic vulnerability, demonstrating that the next-generation JAK inhibitor pacritinib can effectively counteract HMGA1-driven malignancy and mitigate resistance to first-generation JAK inhibitors, thus providing a rational therapeutic avenue for this high-risk patient cohort.

HMGA1’s role as a “molecular rheostat” or “epigenetic fulcrum” in MPN evolution is underscored by its capacity to integrate diverse oncogenic inputs. As an architectural chromatin protein, HMGA1 modulates higher-order chromatin structure and gene accessibility, thereby globally influencing transcriptional landscapes^16,18,31–33^. Our integrated multi-omics data, particularly CUT&Tag and ATAC-seq, compellingly demonstrate that HMGA1 directly occupies and maintains open chromatin at the promoters and enhancers of key G1/S phase regulators (*E2F1*, *CCNE1*/*2*, *CDK2*) and G2/M drivers (*CCNB1*/*2*, *CDC2*), thereby locking-open these critical cell cycle control elements and facilitating the binding of other pro-proliferative transcription factors like AP-1 and ETS. This direct and potent control over cell cycle machinery underscores the significance of HMGA1’s upregulation during MPN progression. Indeed, initial driver mutations, such as *JAK2*^V617F^, induce a STAT5/NF-κB-mediated upregulation of HMGA1, priming key cell-cycle control elements^21,22^. The subsequent acquisition of cooperating lesions, notably dominant-negative or multi-hit *TP53* lesions prevalent in approximately 20–30% of post-MPN AML^34^, then fully unleashes HMGA1’s oncogenic functions. *TP53* mutations are highly significant in this context, being prevalent in 25-50% of post-MPN AML cases and frequently associated with aggressive clonal expansion and poor prognosis, especially when biallelic or compound heterozygous^6^. Murine models have further shown that *Trp53* inactivation in a *JAK2*^V617F^ background can enhance leukemogenic potential and alter therapeutic responses, such as to interferon^7^. Our findings that dominant-negative *TP53* mutations synergize with *JAK2*^V617F^ to maximally induce HMGA1 align with this understanding, suggesting HMGA1 as a key mediator of *TP53*-mutant driven aggressiveness. Our findings that dominant-negative *TP53* mutations synergize with *JAK2^V617F^* to maximally induce HMGA1 align with this understanding, suggesting HMGA1 as a key mediator of *TP53*-mutant driven aggressiveness. This amplified HMGA1 activity likely facilitates the recruitment of transcriptional co-activator complexes to critical genomic loci, thereby potentiating E2F- and MYC-driven proliferation and antagonizing myeloid differentiation. This mechanistic framework offers a cogent explanation for the aggressive clonal dominance and therapeutic refractoriness observed in *TP53*-mutant sAML and is consistent with accelerated disease phenotypes in *Trp53*^R172H^;*Jak2*^V617F^ murine models^27^. Notably, HMGA1’s oncogenic drive extends beyond *TP53*-mutant contexts, implying its function as a convergent downstream effector for a spectrum of high-risk mutations such as *ASXL1*, *SRSF2*, *IDH1*/*2*, *EZH2* that collectively define the complex pathobiology of sAML^4^. Future investigations into whether *Hmga1* ablation can impede disease progression in more complex, compound-mutant murine models as *JAK2*^V617F^ plus *ASXL1*^−/−^ would further solidify the rationale for HMGA1-directed therapies.

The clinical actionability of HMGA1 as a biomarker is a salient translational outcome of this research. HMGA1 IHC, a robust and widely deployable laboratory assay, demonstrated superior predictive accuracy (AUC 0.96) for sAML transformation, surpassing conventional morphological and immunophenotypic markers (CD34, CD117) which often lack early sensitivity or broad applicability across the malignant clone. The challenge of predicting MPN transformation is underscored by the complex and dynamic clonal evolution that characterizes disease progression, often involving the acquisition of multiple new driver mutations^5^. HMGA1 expression dynamically escalates with disease stage, precedes overt blast crisis, and is ubiquitously maintained across diverse leukemic populations, rendering it a more reliable harbinger of impending transformation than current risk models like MIPSS-70. This aligns with the broader trend in hematology towards incorporating molecular markers for more precise risk stratification, as seen in evolving WHO and ICC classifications^35^. Our finding that HMGA1-high status identified a significant cohort of chronic-phase MPN patients who rapidly progressed to sAML within 12 months, while HMGA1-low patients remained stable, underscores its potential to refine risk stratification. Integrating HMGA1 assessment into clinical algorithms could enable proactive identification of high-risk individuals for intensified surveillance, earlier consideration of definitive interventions such as allo-HSCT^36^, or enrollment in trials of interceptive therapies. Non-invasive monitoring of HMGA1 via liquid biopsies may further enhance its clinical utility. Therapeutically, this study positions HMGA1 as a critical determinant of JAK inhibitor response and validates pacritinib as a promising strategy for HMGA1-high sAML, an area with significant unmet need for nevol agents^1^. We establish that HMGA1 overexpression confers resistance to certain JAK inhibitors (ruxolitinib, fedratinib), largely by sustaining E2F/G2-M transcriptional programs. The persistence of JAK2-mutated clones despite first-generation JAK inhibitor therapy is a well-recognized clinical challenge, often mediated by mechanisms such as activation of alternative signaling pathways like ERK^15^. This therapeutic challenge is compounded in HMGA1-high disease, as HMGA1-driven intrinsic malignant cell proliferation and drug resistance would likely render therapies insufficient if their efficacy relies significantly on modulating the microenvironment or non-malignant cell signaling, a mechanism partly attributed to ruxolitinib^37^. Pacritinib, a next-generation JAK2/FLT3/IRAK1 inhibitor, not only retains efficacy in HMGA1-high milieus but also durably suppresses these oncogenic networks^21^. This capacity likely stems from pacritinib’s broader kinase inhibitory profile, including FLT3, IRAK1, and CSF-1R^30,38^, which may exert synergistic anti-leukemic effects in the genetically heterogeneous context of sAML. Even if HMGA1 maintains an open chromatin state at key proliferative loci, pacritinib’s multi-targeted inhibition effectively sterilizes this prepared landscape by cutting off the upstream activating signals required by the transcription factors and signaling cascades that would otherwise exploit these HMGA1-primed sites. The adaptive resistance observed with prolonged ruxolitinib exposure characterized by rebound of E2F1/cyclin B2 and diminished γ-H2AX signaling is effectively circumvented by pacritinib, which sustains DNA damage signaling and cell cycle arrest. This is consistent with its potential modulation of the STAT3/miR-21 axis^39^, highlighting its multifaceted action. Our compelling in vivo data confirm that pacritinib markedly reduces tumor burden and prolongs survival in HMGA1-OE xenografts, effectively neutralizing HMGA1’s adverse impact. These findings advocate for an HMGA1-informed therapeutic algorithm that while ruxolitinib may suffice for HMGA1-low/stable disease, escalating HMGA1 should prompt consideration for pacritinib and evaluation for allo-HSCT.

Future investigations should prioritize the prospective validation of HMGA1 as a predictive biomarker and pacritinib’s efficacy in HMGA1-high MPN/sAML through well-designed, multi-center clinical trials incorporating standardized HMGA1 diagnostics and correlative studies. Exploration of pacritinib-based combination strategies is also critical; its induction of DNA damage suggests synergy with DDR inhibitors such as PARP/ATR inhibitors or BCL-2 antagonists like venetoclax. Given HMGA1’s potential immunomodulatory roles, combinations with immunotherapy also merit investigation. Furthermore, despite the challenges, advancing direct HMGA1-targeting modalities such as small molecule inhibitors, antisense oligonucleotides (ASOs), or proteolysis-targeting chimeras (PROTACs), remains a significant long-term objective.

This study has limitations. Formal shRNA rescue experiments were not performed, precluding definitive exclusion of off-target effects for all knockdown phenotypes. While our clinical cohort is substantial, validation in larger, prospective multi-center cohorts is essential to unequivocally establish HMGA1’s prognostic and predictive utility, particularly for the sAML subset, and to mitigate potential biases from retrospective analyses. Animal models, though informative, incompletely recapitulate human disease complexity; future work with sophisticated patient-derived and humanized models is warranted. Standardization of HMGA1 IHC across laboratories will be crucial for its clinical translation. Finally, the long-term tolerability and potential off-target effects of pacritinib, as a multi-kinase inhibitor, require continued evaluation, although its existing approval provides a foundational safety profile. These limitations delineate clear avenues for future research to refine and extend our conclusions.

In summation, this investigation delineates HMGA1 as a central protagonist in the progression of MPN to sAML, serving as an early warning biomarker, a driver of malignant transformation and therapeutic resistance, and a druggable vulnerability. HMGA1 expression analysis by IHC offers a pragmatic and powerful tool for enhanced risk stratification and timely therapeutic intervention in high-risk MPN. Pacritinib, a clinically available agent, demonstrates compelling activity against HMGA1-driven oncogenic programs and circumvents resistance to first-generation JAK inhibitors, heralding a promising, readily translatable therapeutic strategy. Prospective clinical trials predicated on HMGA1 status are now essential to formally integrate these findings into clinical practice and to explore pacritinib-based combination regimens, with the ultimate aim of improving outcomes for patients confronting this aggressive leukemic transformation.

## Authors’ contributions

Can Yang, Yajun Li, Xuemei Tang, Qingyun Zhang performed experiments. Guan Ming Zhiyuan Wu, Gusheng Tang, Yu Zeng, Wenshu Zheng, and Zhen Huang provided technical and clinical support. Kun Chen, Jianfei Fu, Gushegn Tang evaluated histopathology. Guan Ming and Zhiyuan Wu designed and supervised the experiments. Guan Ming, Can Yang and Zhiyuan Wu wrote the manuscript. Maxiao Tong, Wenjiao Qin, Wenhao Zhang and Jianfei Fu contributed to sample collection. Can Yang and Yajun Li analyzed data. All authors read and approved of the manuscript.

## Acknowledgements

This study was funded by National Natural Science Foundation of China (grant number. 82472369, 82272415 and 81871728), the Innovation Group Project of Shanghai Municipal Health Commission (grant number 2019CXJQ03), Shanghai Municipal Health Commission (grant number 2022YQ045), and Shanghai “Rising Stars of Medical Talents”–Clinical Laboratory Practitioner Program (grant number 2022-065). We thank Shanghai Model Organisms Center, Inc. for offering *Jak2^V617F^* mice, M-NSG mice and identification assistance. We thank the PPL for plasmid construction. We thank the OE Biotech Co., Ltd. for RNA sequencing and analysis. We thank the OE Biotech Co., Ltd. for RNA sequencing, CUT&Tag and analysis. We thank the Shanghai Dache Biotechnology Co., Ltd for ATAC-seq and analysis. We thank the NCBI GEO and OHSU BeatAML for offering public datasets.

---

**Extended Data Fig. 1.**
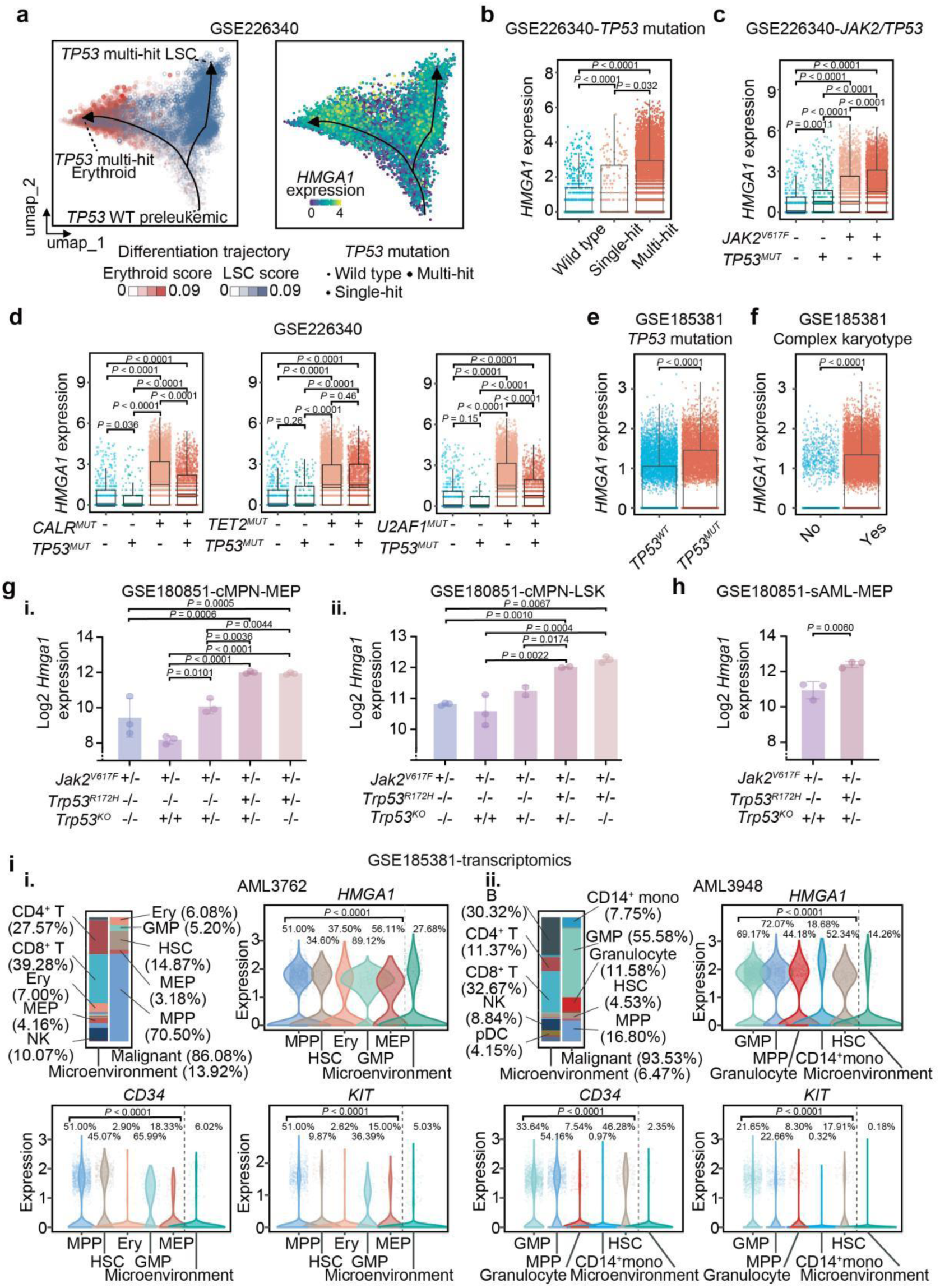
Transcriptomic landscape of HMGA1 expression in MPN-sAML at singel-cell resolution. (a) UMAP visualization of 9,191 Lin^-^CD34^+^ cells from 14 sAML patient samples (GSE226340), showing pseudotime trajectories. Cells are colored by (left to right): differentiation trajectory score (LSC-like vs. erythroid), *HMGA1* expression level, and *TP53* mutational status (wild type, single-hit, multi-hit). Arrows indicate inferred differentiation direction. (b) Violin plots showing *HMGA1* transcript expression in CD34^+^ HSPCs from sAML patients (GSE226340), stratified by *TP53* mutational status (wild type, single-hit, multi-hit). (c) Violin plots showing *HMGA1* transcript expression in CD34^+^ HSPCs from sAML patients (GSE226340), stratified by specific combinations of *JAK2* (wild-type or V617F) and *TP53* (wild-type or mutated) genotype. (d) *HMGA1* expression in CD34^+^ HSPCs from sAML patients (GSE226340) stratified by combinations of *TP53* genotype (wild-type or mutated) with other co-occurring gene mutations (*CALR*, *TET2*, *U2AF1*) versus *TP53* mutation alone. (e) *HMGA1* expression in BMMCs from sAML patients (GSE185381) stratified by *TP53* mutational status (wild type, n = 2 vs. mutated, n = 1). (f) *HMGA1* expression in BMMC from patients with sAML (GSE185381) stratified by karyotype complexity (non-complex karyotype, n = 1 vs. complex karyotype, n = 2) (g) *Hmga1* transcript expression in murine hematopoietic progenitors from various *Jak2*/*Trp53* mutant mouse models (GSE180851). (i) Megakaryocyte-erythorid progenitors (MEP) from *JAK2*^V617F^ (*J^VF^*) mice versus *J^VF^* mice with different *Trp53* genotypes (*Trp53^-/-^*, *Trp53^+/-^*, *Trp53^R172H/-^* or *Trp53^R172H/+^*) (n = 3 per group). (ii) Lin-Sca-1^+^c-kit^+^ (LSK) stem cells from the same genotypes. (h) *HMGA1* expression in MEPs from mice at sAML stage (*Jak2*^V617F^*Trp53^-/-^*, n = 3; *Jak2*^V617F^*Trp53^R172H/-^*, n = 3) (GSE180851). (i) Cellular composition and marker expression in representative sAML cases from GSE185381. (i) Stacking bar charts showing the percentage of annotated cell types in two sAML cases (AML3762, AML3948). (ii) Violin plots depicting *HMGA1*, *CD34*, and *KIT* transcript expression in the top 5 most abundant malignant cell types and microenvironment (non-malignant) cells for each patient. For panels b-h, statistical significance was assessed by Kruskal-Wallis test with Benjamini-Hochberg (BH) correction (b-d), Wilcoxon rank-sum test with BH correction (e,f), likelihood ratio test (g), or Wald test (h) as appropriate.

**Extended Data Fig. 2.**
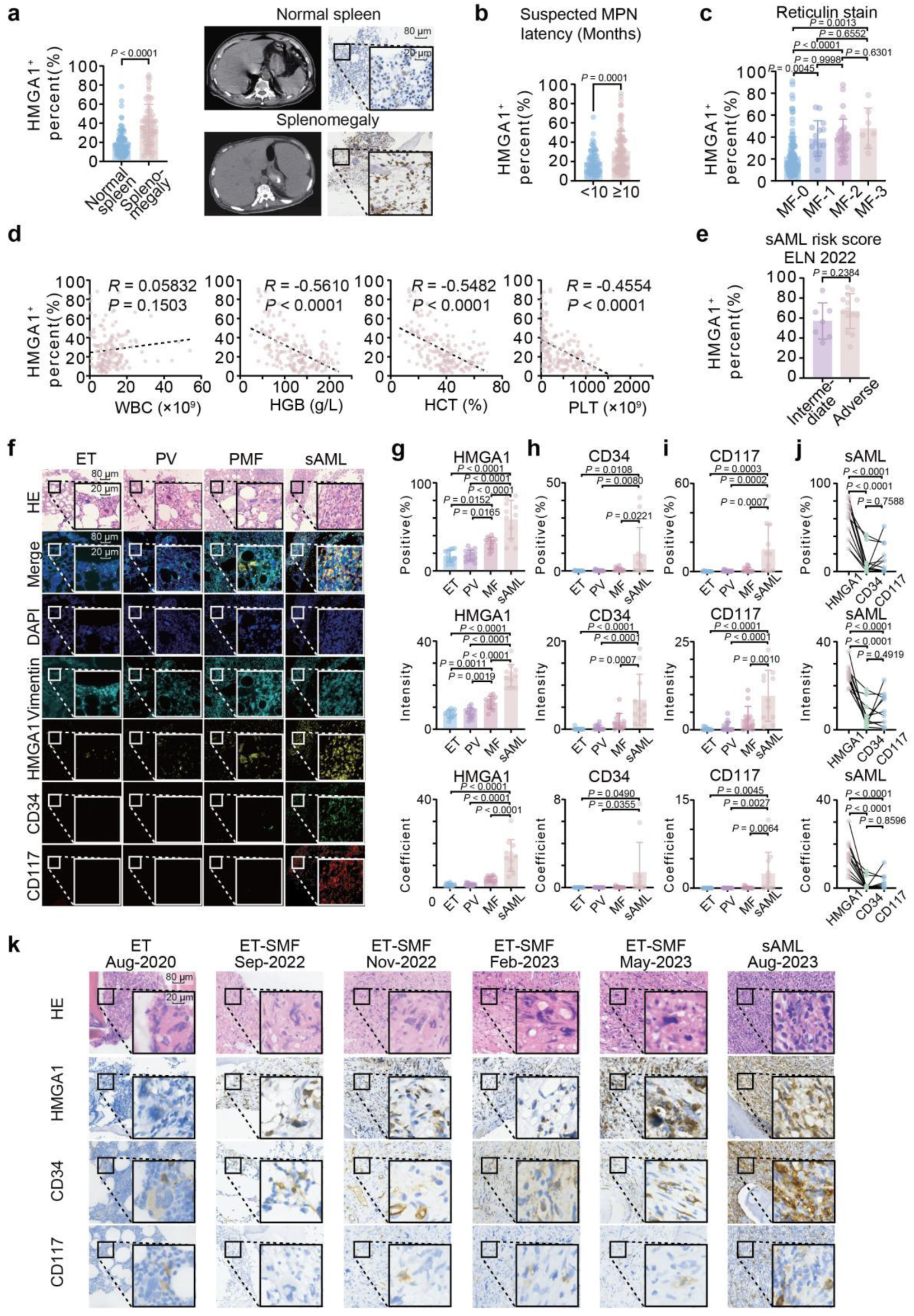
Clinical and pathological correlation of HMGA1 expression in MPN and sAML. (a) HMGA1 IHC score in MPN patients with (n = 95) versus without (n = 67) splenomegaly. Representative IHC images of normal spleen and splenomegaly are shown. Data are mean ± SD. Two-sample *t*-test. Scale bars: top, 80 µm; bottom, 20 µm. (b) HMGA1 IHC scores in MPN patients stratified by the latency period (in months) between initial suspicion of MPN and bone marrow biopsy (< 10 months, n = 66; ≥10 months, n = 90). Data are mean ± SD. Two-sample *t*-test. (c) HMGA1 IHC score in MPN patients stratified by bone marrow reticulin fibrosis grades (MF-0, n = 53; MF-1, n = 33; MF-2, n = 32; MF-3, n = 13). Data are mean ± SD. One-way ANOVA with Tukey’s post-hoc test. (d) Correlation plots of HMGA1 IHC scores versus peripheral blood parameters: white blood cell coutn (WBC), hemoglobin (HGB), hematocrit (HCT), and platelet count (PLT) in the total MPN cohort (n = 162). Pearson correlation coefficient (*r*) and *P*-value are indicated. (e) HMGA1 IHC score in sAML patients (n = 21) stratified by European Leukemia Net (ELN) 2022 risk categories (Intermediate vs. Adverse). Data are mean ± SD. Two-sample *t*-test. (f) Representative multiplex immunofluorescence (mIF) images of bone marrow sections from primary MF and sAML patients, co-stained for HMGA1 (yellow) with CD117 (red), CD34 (green). Nuclei are counterstained with DAPI (blue). HMGA1 is highly expressed in megakaryocytes in PMF and in blasts in sAML. Scale bar: 20 µm. (g-j) Quantification of positive %, intensity and a composite coefficient (positive % × intensity) for (g) HMGA1, (h) CD34, (i) CD117, across different MPN subtypes (ET, n = 12; PV, n = 15; MF, n = 12) and, sAML (n = 11) and (j) three marker in sAML (n = 11). Data are mean ± SD. One-way ANOVA with Tukey’s post-hoc test. (k) Serial H&E and IHC (HMGA1, CD34, CD117) staining of bone marrow biopsies from a representative patient progressing from ET (August 2020) thourgh secondary MF (sMF; September 2022, November 2022, Februrary 2023, May2023) to sAML (August 2023). Scale bars: top, 80 µm; bottom, 20 µm.

**Extended Data Fig. 3.**
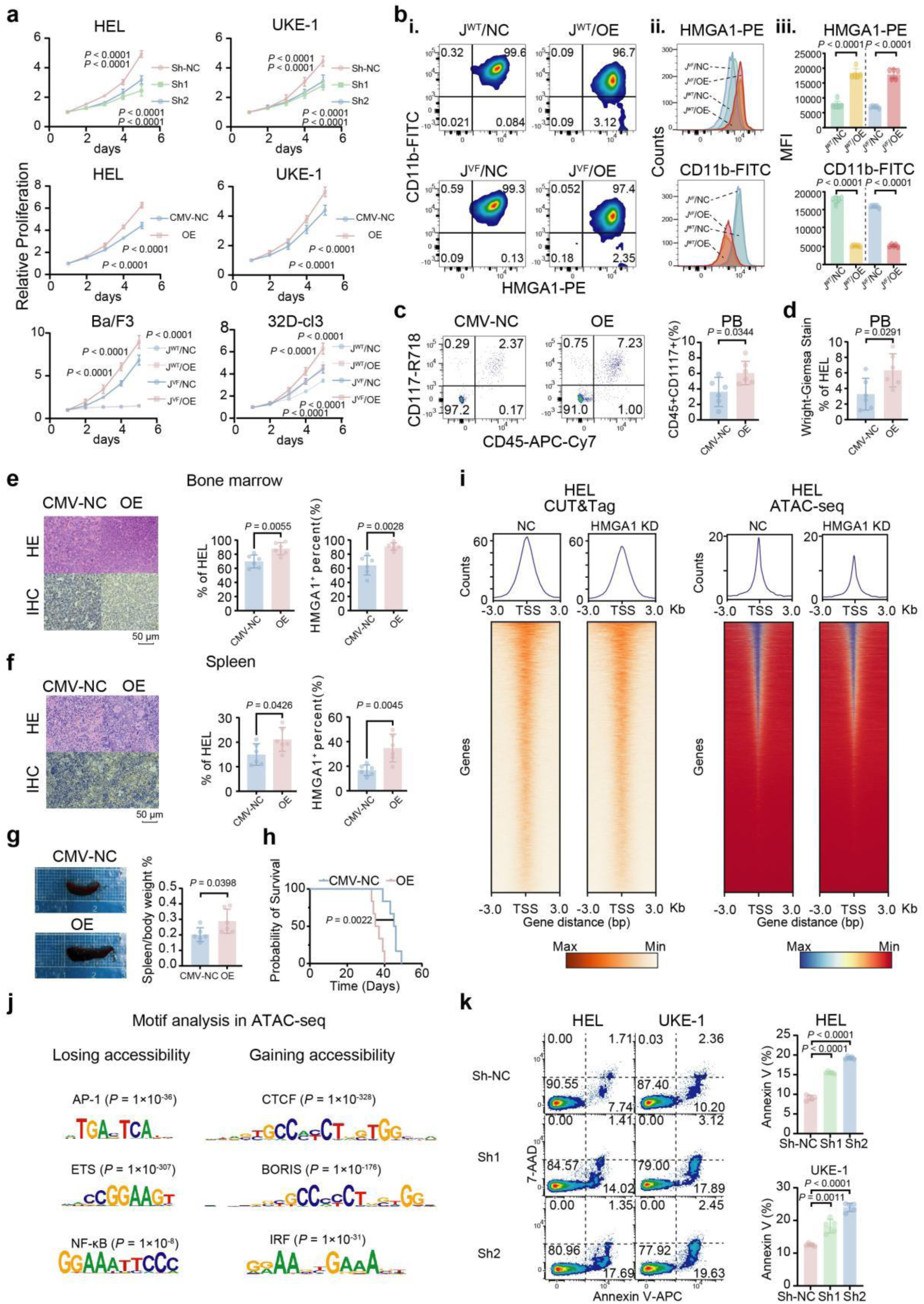
Functional impact of HMGA1 on proliferation, differentiation, and leukemic progression. (a) Relative proliferation curves of human (HEL, UKE-1) and murine (Ba/F3, 32D-cl3 transduced with *Jak2* wild-type or *Jak2*^V617F^) cell lines following HMGA1/Hmga1 overexpression (OE) or shRNA-meidated knockdown (sh1, sh2) compared to respective controls (CMV-NC or sh-NC)NC.) 32D-cl3 cells were cultured with IL-3. Data are mean ± SD. (n = 5 per group). Two-way ANOVA. (b) Flow cytometric analysis of CD11b expression on 32D-cl3 cells transduced with *Jak2* wild-type (J^WT^) or *Jak2*^V617F^(J^VF^), and co-transduced with control vector (NC) or HMGA1 overexpression (OE), following G-CSF (100 ng/mL) induced differentiation. (i) Representative histograms of CD11b-FITC fluorescence. (ii) Quantification of HMGA1-PE mean fluorescence intensity (MFI). (iii) Quantification of CD11b-FITC MFI (n = 5 per group). Data are mean ± SD. Two-sample *t*-test. (c) Quantification of human CD45^+^CD117^+^ HEL cells in peripheral blood of NSG mice at day 35 post-transplant, comparing HMGA1-OE versus vector control (CMV-NC) groups (n=6 per group). Data are mean ± SD. Two-sample *t*-test. (d) Wright-Giemsa stained peripheral blood smears from NSG mice engrafted with HMGA1-OE or CMV-NC HEL cells at day 35. Quantification of HEL cells (% of total nucleated cells) is shown (n = 6 per group). Data are mean ± SD. Two-sample *t*-test. (e-f) Representative H&E and HMGA1 IHC staining (left panels of e and f, respectively) and quantification of HMGA1-positive cells (%) (right panels fo e and f, respectively) in (e) femur bone marrow and (f) spleen sections from NSG mice engrafted with HMGA1-OE or CMV-NC HEL cells. Scale bars: 50 µm. Data are mean ± SD. Two-sample *t*-test. (g) Representative images of spleens (left) and relative spleen weights (spleen weight/body weight %, right) from NSG mice at day 35 post-engraftment with HMGA1-OE or CMV-NC HEL cells (n = 6 per group). Data are mean ± SD. Two-sample *t*-test. (h) Kaplan-Meier survival curves for NSG mice injected with HMGA1-OE ro CMV-NC HEL cells (n = 6 per group). Median survival times are indicated. Log-rank (Mantel-Cox) test. (i) Heatmaps showing HMGA1 binding intensity (CUT&Tag, left) and chromatin accessibility (ATAC-seq, right) centered on transcription start site (TSS ± 3kb) for genes in HEL cells transduced with shNC or shHMGA1. Color scale indicates normalized read counts (Max/Min normalized). (j) Top de novo motifs identified by HOMER analysis within ATAC-seq peak regions that either lose accessibility (left) or gain accessibility (right) upon HMGA1 knockdown in HEL cells. *P*-value for motif enrichment are indicated. (k) Quantification of apoptosis by Annexin V-APC/7-AAD staining and flow cytometry in HEL and UKE-1 cells after transduction with shNC or HMGA1 shRNAs (sh1, sh2). Representative flow cytometry plots are shown. Data are mean ± SD. (n = 5 per group). One-way ANOVA with Tukey’s post-hoc test.

**Extended Data Fig. 4.**
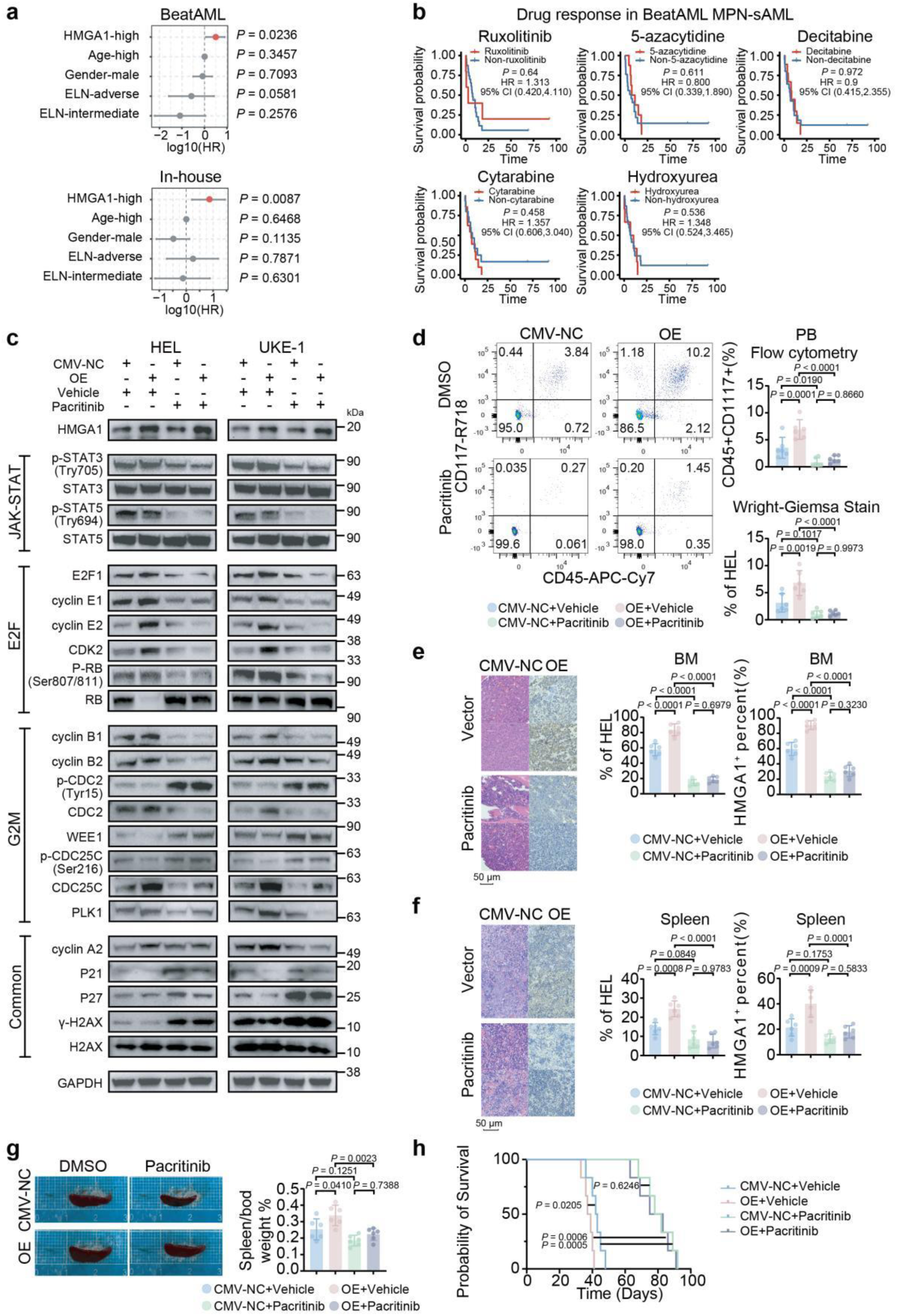
Pacritinib overcomes HMGA1-mediated resistance in MPN-sAML models. (a) Forest plot illustrating multivariable Cox proportional hazards analysis of overall survival (OS) in the OHSU BeatAML cohort (n = 31) and the in-house MPN-sAML cohort (n = 21). Harzard ratio (HR) and 95% confidence intervals (CI) are shown for HMGA1 epxression (high vs. low), age, gender and ELN 2022 risk category. (b) Kaplan-Meier OS curves for MPN-sAML patients from the OHSU BeatAML cohort stratified by specific therapies received (ruxolitinib, 5-azacytidine, decitabine, cytarabine,and hydroxyurea). Log-rank (Mantel-Cox) test. (c) Western blot analysis key proteins in JAK-STAT, E2F, and G2/M pathways in HEL and UKE-1 cells with vector control (CMV-NC) or HMGA1 overexpression (OE), following 4-hour treatment with vehicle (DMSO) or pacritinib (500nM). (d) Quantification of human HEL cells in peripheral blood of NSG mice at day 35 post-transplantation by flow cytometric analysis (CD45^+^CD117^+^) or Writght_Giemsa smear, from mice engrafted with vector control (CMV-NC) or HMGA1-OE HEL cells and treated with vehicle or pacritinib (n=6 mice/group). Data are mean ± SD. One-way ANOVA. (e-f) Pacritinib reduces leukemic infiltration and HMGA1 expression in bone marrow and spleen of HMGA1-OE xenografted mice. Representative IHC staining for HMGA1 and quantification of human HEL cell engraftment (% of HEL cells) and HMGA1-positive cells (%) in (e) bone marrow and (f) spleen sections. NSG mice (n=6 per group) were engrafted with luciferase-expressing HEL cells transduced with CMV-NC or HMGA1-OE constructs, and subsequently treated with vehicle or pacritinib (100 mg/kg, BID) for 14 days. Scale bars for IHC images, 50 µm. Bar graphs depict mean ± SD. One-way ANOVA with Tukey’s post-hoc test. (g) Representative images of spleens and relative spleen weights (spleen weight/body weight %) from NSG mice engrafted with CMV-NC or HMGA1-OE HEL cells, treated with vehicle or pacritinib. One-way ANOVA with Tukey’s post-hoc test. (h) Kaplan–Meier survival curves for NSG mice engrafted with vector control HEL cells (CMV-NC) or HMGA1-OE HEL cells, treated with vehicle or pacritinib (n=6 per group). Log-rank (Mantel-Cox) test.

## Supplementary Information

### Supplementary Methods

#### 1. Patient Cohort and Sample Processing

##### 1.1. Ethical Approval and Consent

The study was conducted in accordance with the Declaration of Helsinki (amended 1996) and received ethical approval from the Institutional Review Boards (IRBs) of Huashan Hospital, Fudan University (Approval No. 2022-1082); Tongji Hospital, Tongji University (Approval No. K-2022-004); and the First Affiliated Hospital of Naval Medical University (Approval No. No. CHEC2022-076). All participants, or their legal guardians, provided written informed consent prior to inclusion in the study.

##### 1.2. Patient Enrollment and Diagnosis

A total of 240 individuals were enrolled between June 2012 and May 2024 at Huashan Hospital, Fudan University; Tongji Hospital, Tongji University; and Changhai Hospital, Navy Military Medical University. This included 162 patients diagnosed with Philadelphia-negative myeloproliferative neoplasms (MPN), encompassing essential thrombocythaemia (ET, n=57), polycythaemia vera (PV, n=33), and myelofibrosis (MF, n=51), or MPN-derived secondary acute myeloid leukemia (sAML, n=21). Additionally, 78 patients initially suspected of having an MPN but ultimately diagnosed with other non-MPN hematological conditions served as a control group.

MPN and sAML diagnoses were established and confirmed according to the 5th edition of the World Health Organization (WHO) classification of hematolymphoid tumors^1^. All diagnoses were subject to central pathology review by at leasttwo independent hematopathologists (K.C., J.F., and G.T.) to ensure diagnostic consistency and accuracy.

##### 1.3. Inclusion and Exclusion Criteria

1. Inclusion criteria:

a. Age between 18 and 90 years.
b. Clinically, morphologically, and molecularly confirmed diagnosis of MPN (PV, ET, MF) or MPN-sAML. MPN-sAML was defined by the presence of ≥ 20% blasts in peripheral blood or bone marrow, and/or extramedullary blast disease, in a patient with a pre-existing or concurrently diagnosed MPN.
c. Adequate hepatic function (e.g., total bilirubin ≤1.5 × upper limit of normal [ULN], AST/ALT ≤2.5 × ULN, unless related to underlying disease), renal function (e.g., creatinine ≤1.5 × ULN or creatinine clearance ≥50 mL/min), and cardiac function (e.g., left ventricular ejection fraction ≥50%) as assessed by standard laboratory tests within 30 days prior to enrollment.
d. Ability to provide written informed consent and comply with study procedures.
2. Exclusion criteria:

a. History of other non-hematologic malignancies within the past 5 years, except for adequately treated non-melanoma skin cancer or carcinoma in situ.
b. Presence of significant comorbidities that, in the investigator’s opinion, could confound the interpretation of study results or pose undue risk to the patient.
c. Current pregnancy or breastfeeding.
d. Known hypersensitivity to bone marrow biopsy procedures or local anesthetic agents. Due to the rarity of sAML (median time from chronic MPN to sAML ∼ 86 months), and the difficulty in accruing a sufficient number of sAML samples for random sampling within the nearly 12-year follow-up period, a convenience sampling strategy was employed for this specific subgroup. Full clinicopathological metadata for all enrolled patients are provided in Supplementary Table S1.

##### 1.4. Sample Collection and Initial Processing

Peripheral blood (PB) and/or bone marrow (BM) aspirate/biopsy samples were collected from healthy donors and patients at the time of diagnosis or during follow-up visits, as clinically indicated.

For PB samples, approximately 5-10 mL was collected into K2-EDTA anticoagulated tubes. For BM aspirates, 2-5 mL was collected into K2-EDTA. BM biopsy specimens were collected into 10% neutral buffered formalin.

Mononuclear cells (PBMCs or BMMCs) were isolated from fresh PB or BM aspirates within 4 hours of collection by Ficoll-Paque PLUS (Cytiva, Cat# 17144002) density gradient centrifugation according to the manufacturer’s instructions. Isolated mononuclear cells were washed twice with phosphate-buffered saline (PBS).

For CD34^+^ cell enrichment, mononuclear cells were resuspended in MACS buffer (PBS supplemented with 0.5% BSA and 2 mM EDTA) and incubated with CD34 MicroBeads (Miltenyi Biotec, Cat# 130-046-702) for 30 minutes at 4°C. CD34^+^ cells were then positively selected using an AutoMACS Pro Separator (Miltenyi Biotec, Cat# 130-092-545) according to the manufacturer’s “possel” program. Purity of CD34^+^ selected cells was routinely assessed by flow cytometry and was typically >90%.

Isolated PBMCs, BMMCs, or enriched CD34^+^ cells were cryopreserved in FBS containing 10% dimethyl sulfoxide (DMSO; Sigma-Aldrich, Cat# D2650) and stored in liquid nitrogen vapor phase until further analysis.

#### 2. Cell Lines and Culture Conditions

##### 2.1. Cell Lines

Human erythroleukemia cell line HEL92.1.7 (ATCC TIB-180), harboring a homozygous *JAK2^V617F^* mutation, and human megakaryoblastic leukemia cell line UKE-1 (Coriell GM23245), also *JAK2^V617F^* positive, were utilized. Murine IL-3-dependent pro-B cell line Ba/F3 (DSMZ ACC 300) and myeloblastic cell line 32D-cl3 (ATCC CRL-11346) were also used. Cell line identities were periodically confirmed by short tandem repeat (STR) profiling. All cell lines were routinely tested for mycoplasma contamination using a PCR-based assay (Lonza, Cat# LT07-318).

##### 2.2. Culture Conditions

All cell lines were cultured in RPMI 1640 medium (Gibco, Cat# 11875093) supplemented with 10% heat-inactivated fetal bovine serum (FBS; Gibco, Cat# 10270106) and 1% penicillin-streptomycin solution (100 U/mL penicillin, 100 µg/mL streptomycin; Gibco, Cat# 15140122). Ba/F3 and 32D-cl3 cells additionally required supplementation with 1 ng/mL recombinant mouse IL-3 (PeproTech, Cat# 213-13) for survival and proliferation, unless otherwise specified for particular experiments (e.g., cytokine withdrawal or differentiation assays). Cells were maintained in a humidified incubator at 37°C with 5% CO2.

#### 3. Molecular Perturbations

##### 3.1. Lentiviral Constructs and Production

Lentiviral vectors for overexpression of human *HMGA1* (NM_145899.3), murine *Hmga1* (NM_016660.3), murin<u>e</u> *Jak2^WT^* (NM_008413.4), or murin<u>e</u> *Jak2^V617F^* were generated by cloning the respective open reading frames (ORFs) into the pLenti-CMV-Puro vector (Addgene plasmid #39481, a gift from Ie-Ming Shih). ORFs were synthesized by GenScript with flanking restriction sites for cloning.

For *HMGA1* knockdown, shRNA sequences targeting human *HMGA1* (sh1: TRCN0000018949; sh2: TRCN0000018951) were obtained from the Sigma-Aldrich MISSION shRNA library and cloned into the pLKO.1-puro vector (Addgene plasmid #8453, a gift from Bob Weinberg). A non-targeting shRNA control (pLKO.1-shSCR; Addgene plasmid #1864) was used.

Lentiviral particles were produced by co-transfecting HEK293T cells (ATCC CRL-3216) with the respective lentiviral plasmid, psPAX2 packaging plasmid (Addgene plasmid #12260, a gift from Didier Trono), and pMD2.G envelope plasmid (Addgene plasmid #12259, a gift from Didier Trono) using Lipofectamine 3000 (Invitrogen, Cat# L3000015). Viral supernatants were harvested at 48 and 72 hours post-transfection, filtered (0.45 µm), and concentrated by ultracentrifugation (25,000 rpm for 2 hours at 4°C) or using Lenti-X Concentrator (Takara Bio, Cat# 631232). Viral titers were determined using a Lenti-X qRT-PCR Titration Kit (Takara Bio, Cat# 631235).

##### 3.2. Transduction and Selection

Target cells (HEL, UKE-1, Ba/F3, 32D-cl3) were transduced with lentiviral particles at a multiplicity of infection (MOI) of 5-10 in the presence of 8 µg/mL polybrene (Sigma-Aldrich, Cat# H9268). After 24-48 hours, transduced cells were selected with 1-2 µg/mL puromycin (Sigma-Aldrich, Cat# P8833) for 7-10 days until non-transduced control cells were completely eliminated. Overexpression or knockdown efficiency was validated by qRT-PCR and Western blotting for *HMGA1*, *Hmga1* or *Jak2* as appropriate.

#### 4. In Vitro Functional Assays

##### 4.1. Cell Viability Assay (CCK-8)

Cells were seeded into 96-well flat-bottom plates at a density of 5 × 10³ (HEL, UKE-1) or 2 × 10⁴ (Ba/F3, 32D-cl3) cells per well in 100 µL of complete culture medium. For drug treatment assays, cells were allowed to stabilize for 4-6 hours before adding serially diluted inhibitors or vehicle control (DMSO, final concentration ≤0.1%). After 72 hours of incubation, 10 µL of Cell Counting Kit-8 solution (CCK8; MCE, Cat# HY-K0301) was added to each well, and plates were incubated for an additional 1-4 hours at 37°C. Absorbance at 450 nm was measured using a Synergy H1 Hybrid Multi-Mode Reader (BioTek). Viability was expressed as a percentage relative to vehicle-treated control cells. Each condition was performed in at least triplicate.

##### 4.2. Apoptosis Assay (Annexin V/7-AAD Staining)

Apoptosis was assessed using the Annexin V-APC/7-AAD Cell Apoptosis Detection Kit (Absin, Cat# abs50008) according to the manufacturer’s protocol. Briefly, 1-10 × 10⁵ cells per condition were harvested, washed twice with ice-cold PBS, and resuspended in 100 µL of 1× binding buffer. Cells were then stained with 5 µL of Annexin V-APC and 10 µL of 7-AAD staining solution for 15 minutes at room temperature in the dark. After staining, 400 µL of 1× binding buffer was added, and samples were analyzed within 1 hour on a BD FACSLyric flow cytometer (BD Biosciences). Data were analyzed using FlowJo software (v10.8.1, BD Life Sciences). Early apoptotic cells (Annexin V^+^/7-AAD^-^), late apoptotic/necrotic cells (Annexin V^+^/7-AAD^+^), and viable cells (Annexin V^-^/7-AAD^-^) were quantified. Each assay was performed in triplicate.

##### 4.3. Cell Cycle Analysis

For cell cycle analysis, the Cell Cycle Detection Kit (Absin, Cat# abs50005) was used. Approximately 1 × 10⁶ cells per sample were harvested, washed with PBS, and fixed by dropwise addition into ice-cold 75% ethanol while vortexing, followed by incubation at - 20°C for at least 2 hours (or overnight at 4°C). Fixed cells were washed with PBS, then resuspended in 500 µL of propidium iodide (PI)/RNase A staining solution and incubated at 37°C in the dark for 30 minutes. DNA content was analyzed using a BD FACSLyric flow cytometer. Cell cycle phase distribution (G0/G1, S, G2/M) was determined using FlowJo software with the Watson (Pragmatic) cell cycle model. Each condition was run in triplicate.

##### 4.4. Cytokine-Stimulated Granulocytic Differentiation

32D-cl3 cells transduced with *Jak2^WT^* or *Jak2^V617F^*, with or without *Hmga1* overexpression, were washed to remove IL-3 and resuspended in RPMI 1640 medium supplemented with 10% FBS, 1% penicillin/streptomycin, and 100 ng/mL recombinant murine Granulocyte-Colony Stimulating Factor (G-CSF; PeproTech, Cat# 250-05). Cells were cultured for up to 14 days, with medium and G-CSF replenished every 3-4 days. Differentiation status was assessed at indicated time points by flow cytometry for the expression of the myeloid differentiation marker CD11b (BD Biosciences, Cat# 557396, clone M1/70) and intracellular HMGA1. All conditions were performed in triplicate.

##### 4.5. Colony-Formation Assay

Cells (HEL, UKE-1, Ba/F3, 32D-cl3 with indicated genetic modifications) were harvested from optimal growth conditions, washed, and resuspended in complete culture medium. One or two thousand viable cells (as determined by Trypan Blue exclusion) were seeded into each well of a 6-well plate containing 2 mL of MethoCult™ GF H4230 (STEMCELL Technologies, Cat# 04230) for human cells or MethoCult™ GF M3134 (STEMCELL Technologies, Cat# 03134) with 1 ng/mL recombinant mouse IL-3 for murine cells, prepared according to the manufacturer’s instructions. Plates were incubated at 37°C in a 5% CO2 humidified atmosphere for 10-14 days. Colonies (defined as aggregates of ≥50 cells) were enumerated under an inverted microscope (Olympus CKX53). Each assay was performed in duplicate or triplicate and repeated at least twice. Representative images of colonies were captured.

#### 5. RNA Sequencing (RNA-seq) and Data Analysis

##### 5.1. RNA Extraction and Quality Control

Total RNA was extracted from 1-5 × 10⁶ cells using TRIzol reagent (ThermoFisher Scientific, Cat# 15596026) according to the manufacturer’s protocol. RNA quantity and purity were assessed using a NanoDrop 2000 spectrophotometer (ThermoFisher Scientific), with A260/A280 ratios typically between 1.8 and 2.1. RNA integrity was evaluated using an Agilent 2100 Bioanalyzer with an RNA 6000 Nano Kit (Agilent Technologies, Cat# 5067-1511); samples with an RNA Integrity Number (RIN) ≥ 7.0 were used for library construction.

##### 5.2. Library Preparation and Sequencing

RNA-seq libraries were constructed from 1 µg of total RNA using the VAHTS Universal V6 RNA-seq Library Prep Kit for Illumina (Vazyme, Cat# NR604-01) following the manufacturer’s instructions. Briefly, mRNA was enriched using oligo (dT) magnetic beads, fragmented, and reverse transcribed into cDNA. cDNA underwent end-repair, A-tailing, adapter ligation, and PCR amplification. Library quality and concentration were assessed using an Agilent 2100 Bioanalyzer and Qubit dsDNA HS Assay Kit (ThermoFisher Scientific, Cat# Q32851). Equimolar amounts of libraries were pooled and sequenced on an Illumina NovaSeq 6000 platform (Illumina Inc.) generating 150 bp paired-end reads. Each experimental condition (e.g., control vs. HMGA1 knockdown) was assessed in biological triplicates.

##### 5.3. RNA-seq Data Analysis

Raw FASTQ reads were subjected to quality control using FastQC (v0.11.9). Adapters and low-quality reads were trimmed using fastp (v0.23.2)^2^ with default parameters. Clean reads were then aligned to the human reference genome (GRCh38/hg38) using HISAT2 (v2.2.1)^3^. Gene expression quantification (read counts per gene) was performed using featureCounts (subread package v2.0.3) ^4^based on Ensembl gene annotations.

Differential gene expression analysis was conducted using DESeq2 (v1.30.1) ^5^in R (v4.2.1). Genes with an adjusted p-value (Benjamini-Hochberg false discovery rate, FDR) < 0.05 and an absolute log2 fold change > 0.5 were considered significantly differentially expressed. Gene Set Enrichment Analysis (GSEA v4.3.2) ^6^was performed using pre-ranked gene lists (ranked by log2 fold change or DESeq2 statistic) against the Hallmark (h.all.v7.5.1.symbols.gmt) and KEGG (c2.cp.kegg.v7.5.1.symbols.gmt) gene sets from the Molecular Signatures Database (MSigDB). An FDR q-value < 0.05 was considered significant for enriched pathways.

#### 6. Assay for Transposase-Accessible Chromatin using sequencing (ATAC-seq)

##### 6.1. Nuclei Isolation and Tagmentation

Approximately 50,000 viable cells per sample were harvested, washed twice with ice-cold 1× PBS (pH 7.4), and resuspended in 50 µL of ice-cold homogenization buffer (10 mM Tris–HCl pH 7.4, 10 mM NaCl, 3 mM MgCl₂, 0.1% IGEPAL CA-630, 0.1% Tween-20, 0.01% Digitonin). Cells were incubated on ice for 3 minutes to lyse plasma membranes and isolate nuclei. Lysates were then washed with 1 mL of ice-cold wash buffer (10 mM Tris-HCl pH 7.4, 10 mM NaCl, 3 mM MgCl₂, 0.1% Tween-20) and centrifuged at 500 × g for 10 minutes at 4°C. The supernatant was discarded, and the nuclear pellet was resuspended in 50 µL of Tn5 transposition reaction mix (25 µL 2× TD Buffer [Illumina, Cat# 20034197], 2.5 µL Tn5 Transposase [Illumina, Cat# 20034197, from Nextera DNA Flex Library Prep Kit], and 22.5 µL nuclease-free water). The transposition reaction was incubated at 37°C for 30 minutes in a thermomixer with gentle shaking (500 rpm). Immediately following transposition, DNA was purified using a MinElute PCR Purification Kit (QIAGEN, Cat# 28004) according to the manufacturer’s instructions, eluting in 21 µL of elution buffer.

##### 6.2. Library Preparation and Sequencing

Transposed DNA fragments were prepared using the TruePrep DNA Library Prep Kit V2 (Vazyme, Cat# TD202) according to the manufacturer’s protocol. PCR products were purified using AMPure XP beads (Beckman Coulter, Cat# A63881) with a double size selection (e.g., 0.5× followed by 1.2× bead ratio) to enrich for fragments between 100 bp and 1000 bp. Library quality and size distribution were assessed using an Agilent 2100 Bioanalyzer with a High Sensitivity DNA Kit (Agilent Technologies, Cat# 5067-4626). Qualified libraries were sequenced on an Illumina NovaSeq 6000 platform generating 150 bp paired-end reads.

##### 6.3. ATAC-seq Data Analysis

Raw FASTQ reads were assessed for quality using FastQC (v0.11.9). Adapters were trimmed using Cutadapt (v3.4)^7^. Clean reads were aligned to the human (GRCh38/hg38) reference genome using Bowtie2 (v2.4.2)^8^. SAM files were converted to BAM, sorted, and indexed using SAMtools (v1.12)^9^. Duplicate reads were removed using Picard MarkDuplicates (v2.25.4). Reads mapping to mitochondrial DNA were removed.

Peak calling on nucleosome-free fragments (typically <150 bp) was performed using MACS2 (v2.2.7.1)^10^. Peak annotation and association with nearby genes were performed using HOMER (v4.11)^11^]. Motif enrichment analysis within accessible regions was also performed using HOMER findMotifsGenome.pl. Differential accessibility analysis between conditions was performed using DESeq2 (v1.30.1) on read counts within consensus peak regions. Peaks with an absolute log2 fold change > 0.5 and an FDR < 0.05 were considered significantly differentially accessible. Visualization of ATAC-seq signal was done using deepTools (v3.5.1) ^12^to generate bigWig files and IGV (Integrative Genomics Viewer v2.11.9)^13^.

#### 7. Cleavage Under Targets and Tagmentation (CUT&Tag)

##### 7.1. CUT&Tag Assay Procedure

CUT&Tag was performed using the Hyperactive In-Situ ChIP Library Prep Kit for Illumina (Vazyme, Cat# TD904) largely following the protocol by Kaya-Okur et al. with some modifications^14^. Briefly, 1 × 10⁵ cells per sample were harvested and washed. Concanavalin A-coated magnetic beads (Bangs Laboratories, Cat# BP531) were activated and used to bind cells. Cells were then permeabilized with wash buffer containing 0.05% Digitonin (Sigma-Aldrich, Cat# D141) and 0.01% IGEPAL CA-630. Cells were incubated with primary antibody against HMGA1 (1:10 dilution; Abcam, Cat# ab129153, rabbit monoclonal) overnight at 4°C in antibody buffer. After washing, cells were incubated with a goat anti-rabbit secondary antibody (1:100; Cell Signaling, Cat# 35401) for 1 hour at room temperature. Subsequently, cells were incubated with protein A-Tn5 (pA-Tn5) transposome complex loaded with Illumina adapters for 1 hour at room temperature. Tagmentation was activated by adding MgCl₂ to a final concentration of 10 mM and incubating at 37°C for 1 hour. The reaction was stopped by adding EDTA, and DNA was released by proteinase K digestion and heating.

##### 7.2. Library Preparation and Sequencing

CUT&Tag DNA fragments were purified using magnetic SPRI beads (Beckman Coulter AMPure XP, Cat# A63881). Libraries were amplified by PCR using indexed primers (NEBNext Multiplex Oligos for Illumina) for 12-15 cycles. Amplified libraries were purified using SPRI beads (double-sided size selection using AMPure XP if necessary) to remove primer dimers and select for appropriate fragment sizes. Library quality and concentration were assessed using an Agilent 2100 Bioanalyzer and Qubit fluorometer. Libraries were sequenced on an Illumina NovaSeq 6000 platform generating 150 bp paired-end reads.

##### 7.3. CUT&Tag Data Analysis

Raw FASTQ reads were processed with fastp (v0.23.2) for quality control and adapter trimming. Clean reads were aligned to the GRCh38/hg38 reference genome using Bowtie2 (v2.4.2). SAM files were converted to BAM, sorted, and indexed. Duplicate reads were removed.

Peak calling was performed using SEACR (Sparse Enrichment Analysis for CUT&RUN; v1.3) in ‘stringent’ mode, an<u>d</u> by selecting the top 0.01 fraction of total signal for normalization^15^. Peaks were annotated to the nearest gene using ChIPseeker (v1.26.2) [PMID: 25765347] in R. Motif analysis of peak regions was performed using MEME Suite (MEME, DREME; v5.3.3)^16^.

Differential peak analysis between HMGA1 knockdown and control groups was performed using MAnorm (v1.3.0)^17^. MAnorm normalizes read counts based on common peaks and uses a Bayesian model to estimate p-values. Significantly differential peaks were defined by FDR < 0.05 and |log2(Fold Change)| > 0.5. Visualization of CUT&Tag signal and peaks was performed using IGV.

#### 8. In Vivo Mouse Models

##### 8.1. *Jak2*^V617F^ Knock-in Mice

To investigate the impact of endogenous *Jak2*^V617F^ expression on Hmga1, Jak2-Flox-V617F/Vav1-Cre-Tg mice (C57BL/6J background; Shanghai Model Organisms Center, Inc., Cat# NM-XA-242440) were utilized. These mice express the *Jak2^V617F^* mutation specifically in hematopoietic cells due to Vav1-Cre-mediated recombination. Age-matched Jak2WT/Vav1-Cre-Tg (wild-type littermates or *Jak2^WT^* controls) mice served as controls. Mice were housed under specific pathogen-free conditions with a 12-hour light/dark cycle and ad libitum access to food and water. At 28 weeks of age, mice (n=5 per genotype) were euthanized by CO2 asphyxiation followed by cervical dislocation. Spleen and femur were collected. Spleens were weighed, and a portion was fixed in 10% neutral buffered formalin for histology. Femurs were also fixed for histological analysis of bone marrow. Peripheral blood was collected via cardiac puncture into EDTA tubes for RNA extraction from PBMCs to assess *Hmga1* transcript levels by RT-qPCR. All animal experiments were conducted in compliance with the guidelines of the Institutional Animal Care and Use Committee (IACUC) of Fudan University.

##### 8.2. Luciferase-Expressing HEL Xenograft Model and Pacritinib Treatment

Six to eight-week-old male NOD.Cg-Prkdc^scid^Il2rg^em1Smoc^ (NSG) mice (Shanghai Model Organisms Center, Inc., Cat# NM-NSG-001) were used for xenograft studies. HEL92.1.7 cells stably transduced with a luciferase reporter gene (pLenti-CMV-Luc2-Blast, Addgene plasmid # 21474, a gift from Eric Campeau) and either control vector (CMV-NC) or HMGA1 overexpression vector (HMGA1-OE) were prepared. Mice were injected intravenously (i.v.) via the tail vein with 3 × 10⁶ viable HEL cells in 100 µL sterile PBS.

###### Disease Monitoring

Tumor engraftment and burden were monitored weekly by bioluminescence imaging (BLI) using an IVIS Spectrum In Vivo Imaging System (PerkinElmer). Mice were injected intraperitoneally (i.p.) with D-luciferin (150 mg/kg; GoldBio, Cat# LUCK-1G) 10-15 minutes before imaging. Total photon flux (photons/second) was quantified from a defined region of interest (ROI) encompassing the whole body.

###### HMGA1 Overexpression Study

Mice engrafted with CMV-NC-Luc HEL cells (n=6) or HMGA1-OE-Luc HEL cells (n=6) were monitored for survival. A separate cohort of mice (n=6 per group) was euthanized at day 35 post-injection for endpoint analyses. Peripheral blood was collected for flow cytometry (human CD45 [BD Biosciences, Cat# 557833, clone 2D1], human CD117 [BD Biosciences, Cat# 752296, clone 104D2]), complete blood counts (CBC), and Wright-Giemsa staining of blood smears. Spleens were excised and weighed. Spleen and femurs were fixed in 10% formalin for histological analysis (H&E staining, IHC for HMGA1). Femurs underwent decalcification in 10% EDTA (pH 7.4) for 7-10 days prior to paraffin embedding.

###### Pacritinib Treatment Study

Twenty-one days post-injection of HMGA1-OE-Luc HEL cells (or CMV-NC-Luc HEL cells for control arm), mice with confirmed engraftment (by BLI) were randomized into treatment groups (n=6 mice/group): vehicle control (0.5% methylcellulose + 0.2% Tween-80 in water) or pacritinib (MedChemExpress, Cat# HY-16379) at 100 mg/kg. Drugs were administered by oral gavage twice daily (BID) for 14 consecutive days. Mice were monitored for body weight changes and signs of toxicity. BLI was performed at baseline (day 21) and at the end of treatment (day 35). A cohort of mice was euthanized at day 35 for endpoint analyses as described above. A separate cohort was monitored for overall survival until humane endpoints were reached (>20% weight loss, hind limb paralysis, moribund state).

#### 9. Histology and Immunostaining

##### 9.1. Hematoxylin and Eosin (H&E) Staining

Formalin-fixed, paraffin-embedded (FFPE) bone marrow (decalcified) and spleen tissues were sectioned at 4 µm. Sections were baked at 60°C for 2 hours, deparaffinized in xylene, and rehydrated through graded ethanol series to water. Slides were stained with Mayer’s hematoxylin (BASO, Cat# BA4025C) for 5-10 minutes, rinsed, differentiated briefly in 0.5% acid-alcohol, and blued in 0.2% lithium carbonate solution. Counterstaining was performed with eosin Y solution (BASO, Cat# BA4025C) for 1-3 minutes. Sections were then dehydrated through graded alcohols, cleared in xylene, and mounted with a neutral resin-based mounting medium. All H&E stains were performed on at least two sections per sample.

##### 9.2. Reticulin Staining

Reticulin staining was performed on FFPE bone marrow biopsy sections using a commercial kit (BASO, Cat# BA4165) according to the manufacturer’s instructions to assess marrow fibrosis. Briefly, sections were deparaffinized, rehydrated, and oxidized with 1% potassium permanganate solution. After bleaching with 2% oxalic acid, sections were sensitized with 2% iron alum solution. Silver impregnation was carried out using ammoniacal silver solution in the dark. Reduction was achieved with 1% formaldehyde solution. Sections were then toned with 0.2% gold chloride, fixed with 5% sodium thiosulfate, dehydrated, cleared, and mounted. Fibrosis was graded according to the European Myelofibrosis Network (EUMNET) criteria (MF-0 to MF-3). Each specimen was stained in triplicate.

##### 9.3. Immunohistochemistry (IHC) and Immunocytochemistry (ICC)

For IHC, 4 µm FFPE sections were deparaffinized and rehydrated. Endogenous peroxidase activity was blocked with 3% hydrogen peroxide in methanol for 15 minutes. Antigen retrieval was performed by heating sections in 10 mM sodium EDTA buffer (pH 9.0) at 95-100°C for 20-30 minutes. Sections were blocked with 5% normal goat serum (Vector Laboratories, Cat# S-1000) for 30 minutes. Primary antibodies (HMGA1, Abcam, Cat# ab129153, 1:5000; CD34, Abcam, Cat# ab81289, 1:200; CD117, Abcam, Cat# ab237794, 1:100) were incubated overnight at 4°C. A biotinylated secondary antibody (Vector Laboratories, Cat# BA-1000, 1:500) was applied for 30 minutes, followed by streptavidin-horseradish peroxidase (HRP) complex (Vector Laboratories, Cat# SA-5704) for 30 minutes. Visualization was achieved using 3,3’-Diaminobenzidine (DAB) substrate-chromogen system (Vector Laboratories, Cat# SK-4105). Sections were counterstained with hematoxylin, dehydrated, cleared, and mounted.

For ICC, HEL and UKE1 cells were cytospun onto slides, air-dried, fixed in 4% paraformaldehyde for 15 minutes, and permeabilized with 0.25% Triton X-100 in PBS for 10 minutes. Staining then proceeded similarly to IHC, starting from the blocking step.

HMGA1 positivity was scored as the percentage of positively stained nuclei in at least three representative high-power fields (HPF, 400× magnification), counting a minimum of 500 cells per sample.

##### 9.4. Multiplex Immunofluorescence (mIF)

FFPE sections were pretreated as for IHC (deparaffinization, rehydration, antigen retrieval). Multiplex staining was performed using the tyramide signal amplification (TSA)-based technology. Briefly, sections were incubated sequentially with primary antibodies: HMGA1 (Abcam, Cat# ab129153, 1:5000), CD34 (Abcam, Cat# ab81289, 1:200), CD117 (Abcam, Cat# ab32363, 1:100), and Vimentin (Abcam, Cat# ab92547, 1:500). After each primary antibody, an HRP-conjugated secondary antibody was applied, followed by an fluorophore (Cy5, SpGold, SpGreen, SpAqua) diluted in amplification buffer. Antibody stripping (e.g., heat-induced epitope retrieval) was performed between each primary antibody-TSA cycle. Nuclei were counterstained with DAPI (1 µg/mL solution). Slides were mounted with ProLong Gold Antifade Mountant (Invitrogen, Cat# P36930).

Images were acquired using a confocal microscope (Leica TCS SP8) or a slide scanner equipped for multiplex fluorescence (Akoya Vectra Polaris). Image analysis was performed using ImageJ/Fiji (v1.8.0) with appropriate plugins. The positivity rate for HMGA1, CD34, and CD117 was expressed as the percentage of cells positive for the respective marker (IF signal co-localizing with DAPI-stained nuclei) in at least three representative fields. Signal intensity was also quantified. A composite coefficient was sometimes calculated by multiplying the positive cell ratio by the mean intensity.

#### 10. Western Blotting (Immunoblotting)

Cells (typically 1-5 × 10⁶) were seeded and subjected to genetic perturbation or drug treatment for the specified duration. After treatment, cells were harvested, washed twice with ice-cold PBS, and lysed on ice for 30 minutes in RIPA buffer (50 mM Tris-HCl pH 7.4, 150 mM NaCl, 1% NP-40, 0.5% sodium deoxycholate, 0.1% SDS) supplemented with protease inhibitor cocktail (Roche, Cat# 11836170001) and phosphatase inhibitor cocktail (Roche, Cat# 04906837001). Lysates were clarified by centrifugation at 14,000 × g for 15 minutes at 4°C. Protein concentrations were determined using the BCA Protein Assay Kit (ThermoFisher Scientific, Cat# 23225).

Equal amounts of protein (20-40 µg per lane) were mixed with Laemmli sample buffer, boiled for 5 minutes, and resolved by SDS-PAGE on 8-15% polyacrylamide gels. Proteins were then transferred to polyvinylidene difluoride (PVDF) membranes (Millipore, Cat# IPVH00010). Membranes were blocked for 1 hour at room temperature in 5% non-fat dry milk or 5% BSA in Tris-buffered saline with 0.1% Tween-20 (TBST). Membranes were incubated with primary antibodies (dilutions and sources in Supplementary Table S3) overnight at 4°C with gentle agitation. After washing three times with TBST, membranes were incubated with HRP-conjugated secondary antibodies (e.g., goat anti-rabbit IgG-HRP, Cell Signaling Technology, Cat# 7074, 1:2000; goat anti-mouse IgG-HRP, Cell Signaling Technology, Cat# 7076, 1:2000) for 1 hour at room temperature. Protein bands were visualized using an enhanced chemiluminescence (ECL) detection system (SuperSigna West Pico PLUS, ThermoFisher Scientific, Cat# 34580) and imaged using a Tanon™ 5200 Multi Imaging System. Band intensities were quantified using ImageJ software where appropriate. β-actin (Proteintech, Cat# 66009-1-Ig, 1:5000), GAPDH (Proteintech, Cat# 60004-1-Ig, 1:5000) or acetyl-tubulin (Proteintech, at# 66200-1-Ig, 1:5000) served as loading controls. Each immunoblot was repeated independently at least twice, and representative blots are shown.

#### 11. Quantitative Reverse Transcription PCR (qRT-PCR)

Total RNA was isolated from cultured cells (0.5-1 × 10⁶) using TRIzol reagent as described in section 5.1. One microgram of total RNA was reverse transcribed into complementary DNA (cDNA) using the PrimeScript™ RT Reagent Kit with gDNA Eraser (Perfect Real Time; Takara Bio, Cat# RR047A) according to the manufacturer’s protocol.

Quantitative PCR was performed using Taq Pro Universal SYBR qPCR Master Mix (Vazyme, Cat# Q712-02) on a QuantStudio 5 Real-Time PCR System (Applied Biosystems). Each reaction (20 µL total volume) contained 10 µL of 2× SYBR Green Master Mix, 0.4 µL of each primer (10 µM stock), 2 µL of cDNA template (diluted 1:5 or 1:10), and nuclease-free water. Thermocycling conditions were: 95°C for 30 seconds, followed by 40 cycles of 95°C for 10 seconds and 60°C for 30 seconds, followed by a melt curve analysis. Relative mRNA expression levels were calculated using the 2-ΔΔCt method, with β-actin (*ACTB* or *Actb*) used as the endogenous reference gene for normalization. All primer sequences used are listed in Supplementary Table S2. All qRT-PCR experiments were performed in triplicate for each sample and repeated at least twice.

#### 12. Bioinformatics Analysis of Publicly Available Datasets

##### 12.1. Single-Cell Multi-Omics Data

###### 12.1.1. GSE185381 (CITE-seq of MPN-sAML and Healthy Controls)^18^

Normalized CITE-seq (RNA and ADT counts) data from BMMCs of 3 sAML patients and 10 healthy controls were downloaded. Provided metadata (cell malignancy status, patient ID, cell subtype annotations, clinical information) were integrated into a Seurat object (Seurat v5.0.1)^19^. Differential gene expression (DGE) between malignant and non-malignant cells, or between disease stages, was performed using the FindMarkers function (Wilcoxon rank-sum test or Kruskal-Wallis test as appropriate), with p-values adjusted using Benjamini-Hochberg (BH) correction (FDR < 0.05 considered significant). Cell type proportions were calculated and visualized. Violin plots for gene/protein expression were generated. Gene Set Enrichment Analysis (GSEA v4.3.2) was performed on pre-ranked gene lists against Hallmark gene sets.

###### 12.1.2. GSE226340 (TARGET-seq of *TP53*-mutant sAML CD34^+^ HSPCs)^20^

Processed TARGET-seq data, including gene expression, mutation profiles (*TP53*, *JAK2*, *CALR* etc.), and cell annotations (malignant cell type, erythroid/LSC scores) for CD34^+^ HSPCs from 14 *TP53*-mutant sAML patients, were obtained. Cluster analysis was performed using SingCellaR (v2.0)^21^. Pseudotime trajectory analysis to model differentiation paths was conducted using Monocle3 (v1.2.9) ^22^with default parameters. DGE along pseudotime or between mutation-defined subclones was assessed using appropriate statistical tests within the respective packages, with BH correction (FDR < 0.05).

###### 12.1.3. GSE221946 (Paired scATAC-seq and snRNA-seq of PMF to sAML transformation) ^23^

Single-cell ATAC-seq (scATAC-seq) and single-nucleus RNA-seq (snRNA-seq) data from a patient at primary myelofibrosis (PMF) stage and subsequent sAML transformation were downloaded. Data were processed and normalized using Seurat (v5.0.1). For scATAC-seq, peaks were called using MACS2 (v2.2.7.1) and differential accessibility between PMF and sAML CD34^+^ cells was assessed using logistic regression models implemented in Seurat or by comparing read counts in consensus peaks with DESeq2. For snRNA-seq, DGE in CD34^+^ cells between stages was performed using FindMarkers (Wilcoxon test, BH correction). *HMGA1* expression was correlated with disease stage and changes in chromatin accessibility.

##### 12.2. Bulk Transcriptome and ATAC-seq Data from MPN Cohorts

###### 12.2.1. GSE189979 (RNA-seq of CD34^+^ PBMCs from MPN subtypes)^24^

RNA-seq data from CD34^+^ PBMCs of patients with ET (n=2), PV (n=1), and MF (n=5) were analyzed. Raw counts files were processed as described in section 5.3. DESeq2 (v1.30.1) was used for normalization and to compare HMGA1 expression across MPN subtypes.

###### 12.2.2. GSE180851 (RNA-seq from *Jak2^V617F^* mouse models)^25^

RNA-seq data from hematopoietic progenitors (MEP, LSK cells) of *Jak2*^V617F^, *Jak2*^V617F^/*Trp53* various genotypes (*Trp53*^-/-^, *Trp53*^+/-^, *Trp53*^R172H/-^, *Trp53*^R172H/+^) mouse models were obtained. Data were processed and normalized using DESeq2. *Hmga1* expression was compared across genotypes using Wald tests within DESeq2 (*FDR* < 0.05).

###### 12.2.3. GSE210253 (RNA-seq of paired MF and sAML samples)^26^

RNA-seq data from 11 paired MF and subsequent sAML patient samples, with associated mutation data (VAFs for driver and high-risk mutations), were analyzed. Data were normalized using DESeq2. *HMGA1* expression changes and VAFs were compared between paired MF and sAML samples using paired Student’s t-tests.

###### 12.2.4. GSE189570 (RNA-seq and ATAC-seq of HMGA1 KD in DAMI cells)^27^

Bulk RNA-seq and ATAC-seq data from the DAMI cell line with and without HMGA1 knockdown were obtained. RNA-seq data were analyzed using DESeq2 for DGE, followed by GSEA. ATAC-seq data were analyzed as described in section 6.3 to identify differentially accessible regions and associated pathways.

##### 12.3. Transcriptome Analysis of JAK2 Inhibitor-Treated HEL Cells GSE229712 ^28^ and GSE190517^29^

RNA-seq data from HEL cells treated with DMSO (control) or various JAK2 inhibitors (ruxolitinib, fedratinib, pacritinib, momelotinib; typically 500 nM) for short (4 or 48 hours) or persistent (long-term, 30 days for ruxolitinib/fedratinib resistance models) durations were downloaded. Raw reads were processed and analyzed using DESeq2 for DGE (*FDR* < 0.05). GSEA was performed against Hallmark gene sets, with a focus on E2F targets and G2/M checkpoint pathways.

#### 13. Drug Response and Survival Analysis in MPN-sAML Patient Cohorts

##### 13.1. OHSU BeatAML Cohort ^30^

Clinical and gene expression (RNA-seq) data from the Oregon Health & Science University (OHSU) BeatAML dataset were accessed via cBioPortal. Patients classified as sAML with a prior MPN diagnosis were included in our MPN-sAML cohort (n=31). Overall survival (OS) was analyzed using Kaplan-Meier methods and log-rank tests, stratified by HMGA1 expression levels (median or quartile cut-offs). Cox proportional hazards models were used to assess the prognostic impact of *HMGA1*, adjusting for age, sex, and treatment. For GSEA based on hazard ratios (HR), genes were ranked by their HR for association with OS, and enrichment analysis was performed (FDR < 0.05). Response rates (CR/CRi) to specific therapies (ruxolitinib, 5-azacytidine, decitabine, cytarabine, hydroxyurea) were compared between HMGA1-high and HMGA1-low groups using Fisher’s exact test or Chi-squared test.

##### 13.2. In-House Patient Cohort

For our in-house cohort of MPN-sAML patients (n=21), HMGA1 protein expression (assessed by IHC score in bone marrow biopsies) was correlated with clinical outcomes, including response to therapy (CR, CRh, CRi according to ELN 2022 AML criteria) and OS. Statistical methods were similar to those used for the BeatAML cohort.

#### 14. In Vitro Drug Sensitivity Assays (IC₅₀ Determination)

Cells (HEL, UKE-1, Ba/F3, 32D-cl3, with or without *HMGA1*/*Jak2* modifications) were seeded in 96-well plates (5 × 10³ to 2 × 10⁴ cells/well). After 24 hours, cells were treated with a range of concentrations (typically 7-10 points, 3-fold or 4-fold serial dilutions) of various inhibitors (ruxolitinib, fedratinib, pacritinib, momelotinib, IFNα, 5-azacytidine, decitabine, cytarabine, venetoclax, hydroxyurea; sources and catalog numbers in Supplementary Table S3) or vehicle control (DMSO ≤0.1%). After 72-96 hours incubation, cell viability was assessed using the CCK-8 assay. Data were normalized to vehicle-treated controls (100% viability). IC_50_ values (half-maximal inhibitory concentration) were calculated by fitting a four-parameter logistic (variable slope) non-linear regression model to the dose-response curves using GraphPad Prism v8.0. Each drug concentration was tested in triplicate, and experiments were repeated at least twice.

#### 15. Generation of Ruxolitinib-Persistent Cell Lines

To establish ruxolitinib-persistent (Rux-P) subclones, parental HEL and UKE-1 cells were continuously cultured in RPMI 1640 medium supplemented with 10% FBS, 1% penicillin/streptomycin, and gradually increasing concentrations of ruxolitinib (MedChemExpress, Cat# HY-50856), starting from 100 nM up to a final concentration of 1 µM over a period of approximately 1 months. Medium containing ruxolitinib was refreshed every 2-3 days. Once cells consistently proliferated in 1 µM ruxolitinib, they were considered ruxolitinib-persistent. These Rux-P cells were then maintained in medium containing 1 µM ruxolitinib. HMGA1 expression in Rux-P cells compared to parental cells was assessed by RT-qPCR and Western blotting.

## Supplementary Figure Legends

**Supplementary Figure 1.**
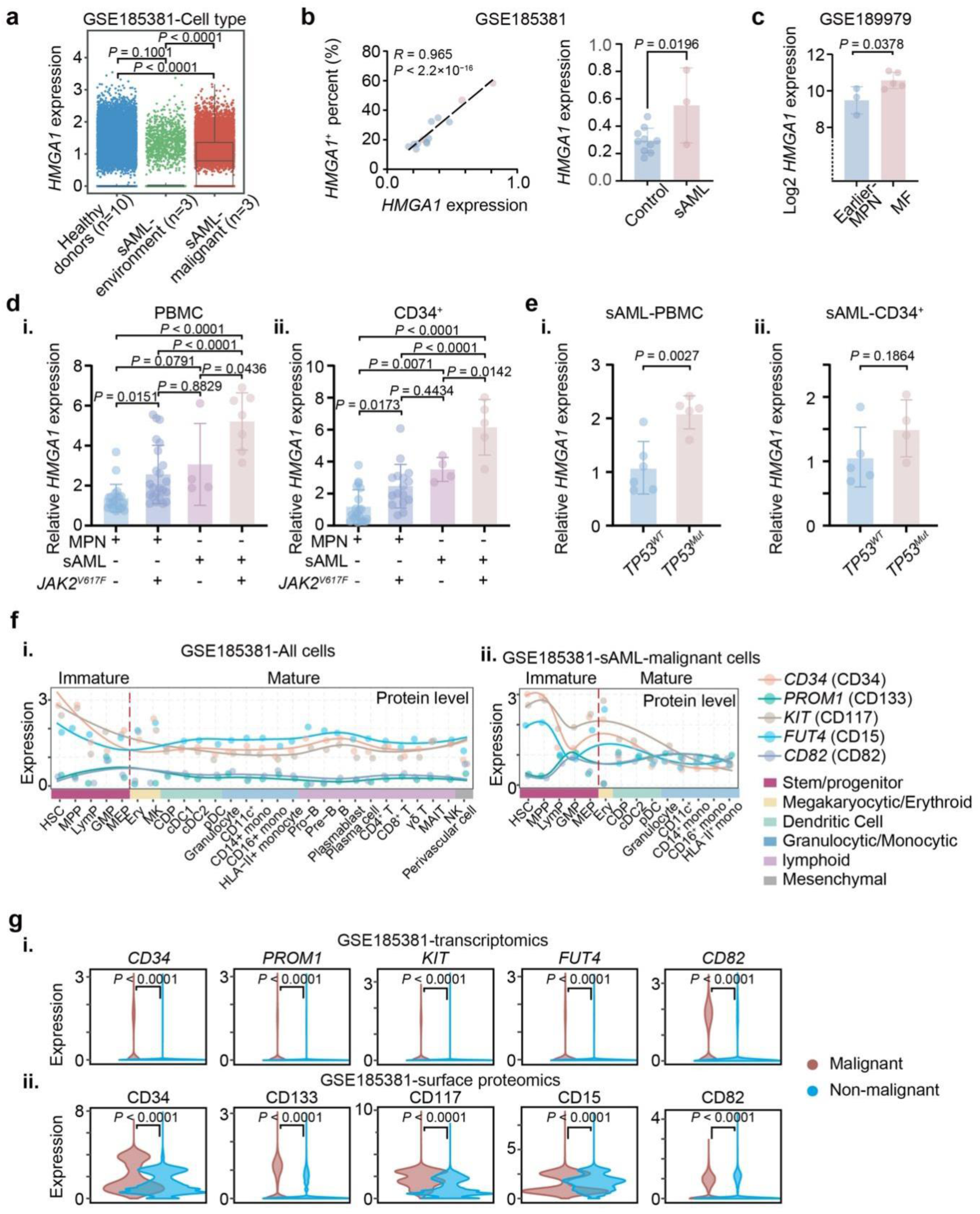
Upregulation of HMGA1 in MPN-sAML at single-cell and bulk transcriptomic levels. (a) Box plots showing HMGA1 expression in bone marrow mononuclear cells (BMMCs) from healthy individuals (n = 10) and sAML patients (n = 3; malignant cells and microenvironment components depicted separately), based on CITE-seq data (GSE185381). Statistical analysis by Kruskal-Wallis test with Benjamini-Hochberg (BH) correction. (b) Left: Correlation between mean *HMGA1* transcript expression and percentage of HMGA1-positive cells from healthy controls and sAML patients (GSE185381). Right: Bar plot comparing mean *HMGA1* expression in total BMMCs between healthy controls (n = 10) and sAML patients (n = 3). Pearson correlation coefficient (*R*) and *P*-value shown. Two-sample *t*-test with BH correction. (c) Log2-transformed *HMGA1* expression in CD34^+^ peripheral blood mononuclear cells (PBMCs) from patients with chronic-phase MPN (ET, n = 2; PV, n = 1; MF, n = 5), derived from dataset GSE189979. Wald test with BH correction. (d) Relative *HMGA1* mRNA expression, determined by qRT-PCR, in (i) PBMCs and (ii) CD34^+^ cells from MPN patients (stratified by JAK2 mutation status: *JAK2* wile-type PBMCs, n = 21; *JAK2*^V617F^ PBMCs, n = 23; *JAK2* wile-type CD34^+^, n = 20; *JAK2*^V617F^ CD34^+^, n = 16) and sAML patients (stratified by *JAK2* mutation status: *JAK2* wile-type sAML PBMCs, n = 4; *JAK2*^V617F^ PBMCs, n = 7; *JAK2* wile-type CD34^+^, n = 4; *JAK2*^V617F^ CD34^+^, n = 5). Data are presented as mean ± SD. One-way ANOVA. (e) Relative *HMGA1* mRNA expression, determined by qRT-PCR, in (i) PBMCs from sAML patients (stratified by *TP53* mutation status: *TP53* wild-type, n = 6; *TP53* mutated, n = 5) and (ii) CD34^+^ cells from sAML patients (*TP53* wild-type, n = 5; *TP53* mutated, n = 4). Data are presented as mean ± SD. Two-sample *t*-test. (f) Expression trajectory plot depicting protein expression of HMGA1 and key stem/progenitor markers (CD34, *PROM1*/CD133, *KIT*/CD117, *FUT4*/CD15, CD82) across diverse hematopoietic cell subsets, ordered by developmental hierarchy, within (i) total BMMCs and (ii) the malignant sAML cell compartment (GSE185381). (g) Violin plots comparing (i) transcript levels (CITE-seq) and (ii) surface protein expression (ADT) of CD34, *PROM1* (CD133), *KIT* (CD117), *FUT4* (CD15), and CD82 between malignant and non-malignant BMMCs from healthy controls (n = 10) and sAML patients (n = 3) (GSE185381). Wald test with BH correction.

**Supplementary Figure 2.**
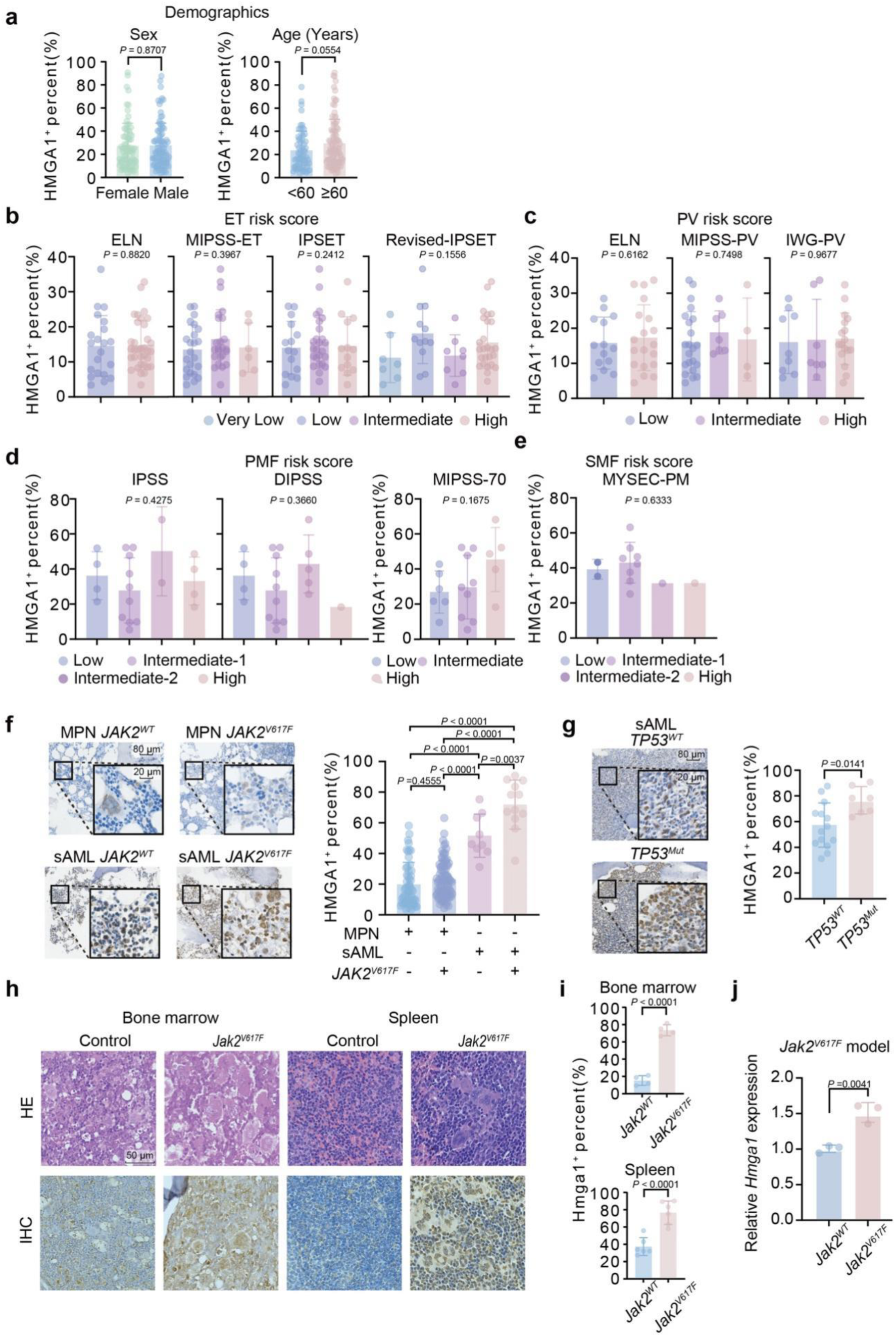
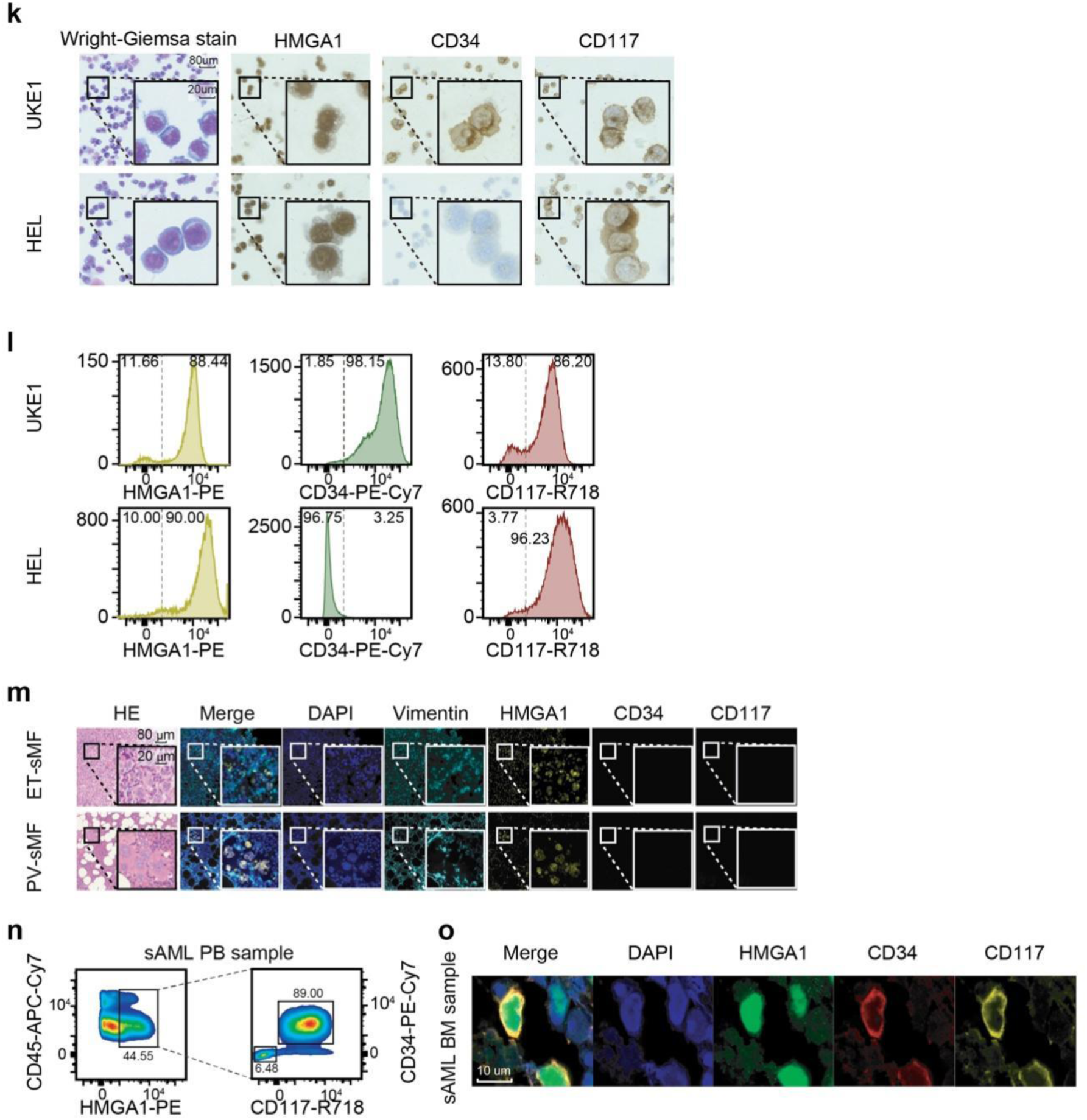
Clinical relevance of HMGA1 expression in MPN and sAML. (a) HMGA1 protein expression (percentage of IHC-positive cells) in bone marrow biopsies from a patient cohort, stratified by sex (female, n = 140; male, n = 67) and age (<60 years, n = 56; ≥60 years, n = 75). Statistical analysis by two-sample t-test. (b-c) Association of HMGA1 protein levels (IHC) with established clinical risk scores in (b) essential thrombocythemia (ET, n = 53) using ELN, MIPSS-ET, IPSET, and Revised-IPSET scoring systems, and (c) polycythemia vera (PV, n = 33) using ELN, MIPSS-PV, and IWG-PV scoring systems. Two-sample *t*-test or one-way ANOVA, as appropriate. (d-e) HMGA1 protein levels (IHC) in (d) primary myelofibrosis (PMF, n = 20) stratified by IPSS, DIPSS, and MIPSS-70 clinical risk scores, and (e) secondary myelofibrosis (sMF, n = 12) stratified by MYSEC-PM score. One-way ANOVA. (f) Differential HMGA1 protein expression (IHC) between MPN (*JAK2* wild-type, n = 53; *JAK2*^V617F^, n = 88) and sAML (*JAK2* wild-type, n = 9; *JAK2*^V617F^, n = 12). Representative IHC images for *JAK2* wild-type MPN and sAML are inset. One-way ANOVA. (g) HMGA1 protein expression (IHC) in sAML patient samples stratified by *TP53* mutation status (*TP53* wild-type, n = 14; *TP53* mutated, n = 7). A representative IHC image for *TP53* wild-type sAML is inset. Two-sample *t*-test. (h-j) Elevated Hmga1 expression in a *Jak2*^V617F^ knock-in MPN mouse model. (h) Representative images of Hematoxylin and Eosin (H&E) staining and IHC for Hmga1 in spleen and bone marrow tissues from *Jak2* wild-type (control) and *Jak2*^V617F^ mice. Scale bars, 50 µm. (i) Quantification of Hmga1 IHC-positive cells (%) in spleen (n = 6 per group) and bone marrow (n = 4 per group). (j) Relative *Hmga1* mRNA levels in PBMCs from *Jak2* wild-type (n = 3) and *Jak2*^V617F^ (n = 3) mice, determined by qRT-PCR. Data are presented as mean ± SD. Two-sample *t*-test. (k-l) Expression profiling of HMGA1, CD34, and CD117 in HEL and UKE-1 MPN cell lines. (k) Representative images of Wright-Giemsa staining (left panels) and immunocytochemical (ICC) staining for HMGA1, CD34, and CD117 (right panels). Scale bars, 80 µm (overview), 20 µm (inset). (l) Flow cytometric analysis of intracellular HMGA1 and surface CD34/CD117 expression. (m) Multiplex immunofluorescence (mIF) demonstrating HMGA1 co-expression with stem/progenitor markers in bone marrow biopsies from MPN patients progressing to sMF (representative PV-sMF and ET-sMF cases). Images show H&E staining alongside fluorescence channels for DAPI (nuclei, blue), Vimentin (cyan), HMGA1 (yellow), CD34 (green), and CD117 (red). Scale bars, 80 µm (overview), 20 µm (inset). (n) Flow cytometric analysis revealing HMGA1, CD34, and CD117 expression within CD45^+^ blasts from an sAML patient’s peripheral blood (PB). (o) Confocal microscopy demonstrating co-localization of HMGA1 (green) with CD34 (red) and CD117 (yellow) in bone marrow (BM) cells from an sAML patient. Nuclei are counterstained with DAPI (blue). Scale bar, 10 µm.

**Supplementary Figure 3.**
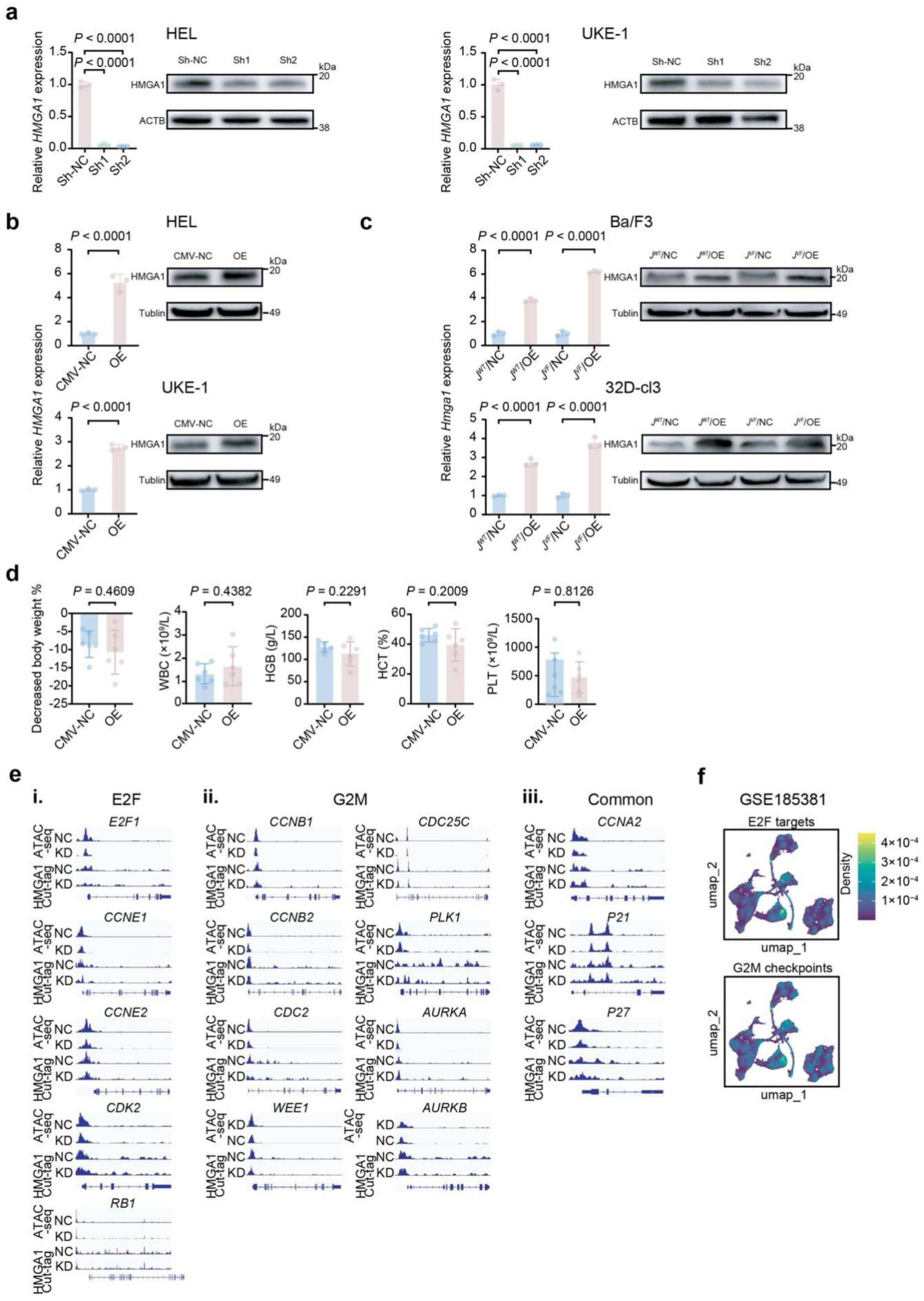
HMGA1 promotes MPN cell proliferation, stemness, and accelerates leukemic progression in vitro and in vivo. (a) Efficient shRNA-mediated knockdown of HMGA1 (sh1, sh2 vs. sh-NC control) in HEL and UKE-1 cells. Left: Relative *HMGA1* mRNA levels by qRT-PCR (mean ± SD, n = 3). Right: Western blot analysis of HMGA1 protein; ACTB served as loading control. (b) Lentiviral-mediated overexpression of HMGA1 (OE vs. CMV-NC control) in HEL and UKE-1 cells. Left: Relative *HMGA1* mRNA levels by qRT-PCR (mean ± SD, n = 3). Right: Western blot analysis of HMGA1 protein; Tubulin served as loading control. (c) Lentiviral-mediated overexpression of Hmga1 (J/OE vs. J/NC control) in murine Ba/F3 (*Jak2* wild type, or *Jak2*^V617F^) and 32D-cl3 (*Jak2* wild type, or *Jak2*^V617F^) cells. Left: Relative *Hmga1* mRNA levels by qRT-PCR (mean ± SD, n = 3). Right: Western blot analysis of Hmga1 protein; Tubulin served as loading control. Statistical analyses for (a-c) by two-sample t-test or one-way ANOVA, as appropriate. (d) HMGA1 overexpression exacerbates disease phenotype in a HEL xenograft model. Hematological parameters (WBC, white blood cell count; HGB, hemoglobin; HCT, hematocrit; PLT, platelet count) in NSG mice engrafted with HEL cells stably expressing control vector (CMV-NC, n = 6) or HMGA1 (OE, n = 6) at 35 days post-transplantation. Data are presented as mean ± SD. Two-sample *t*-test. (e) HMGA1 knockdown alters chromatin accessibility and HMGA1 binding at key cell cycle regulatory gene loci. Integrative Genomics Viewer (IGV) snapshots displaying ATAC-seq and HMGA1 CUT&Tag signals at representative E2F target genes (*E2F1*, *CCNE1*, *CCNE2*, *CDK2*, *RB1*), G2M checkpoint genes (*CCNB1*, *CCNB2*, *CDC2*, *WEE1*, *CDC25C*, *PLK1*, *AURKA*, *AURKB*), and common cell cycle regulators (*CCNA2*, *CDKN1A*/*p21*, *CDKN1B*/*p27*) in HEL cells following control (NC) versus HMGA1 knockdown (KD). (f) Enhanced E2F target and G2M checkpoint gene signatures in sAML patient cells. UMAP projections of single-cell CITE-seq data (GSE185381) from control and sAML patients, with cells colored by enrichment scores for E2F target and G2M checkpoint gene sets. Corresponding density plots illustrate score distributions.

**Supplementary Figure 4.**
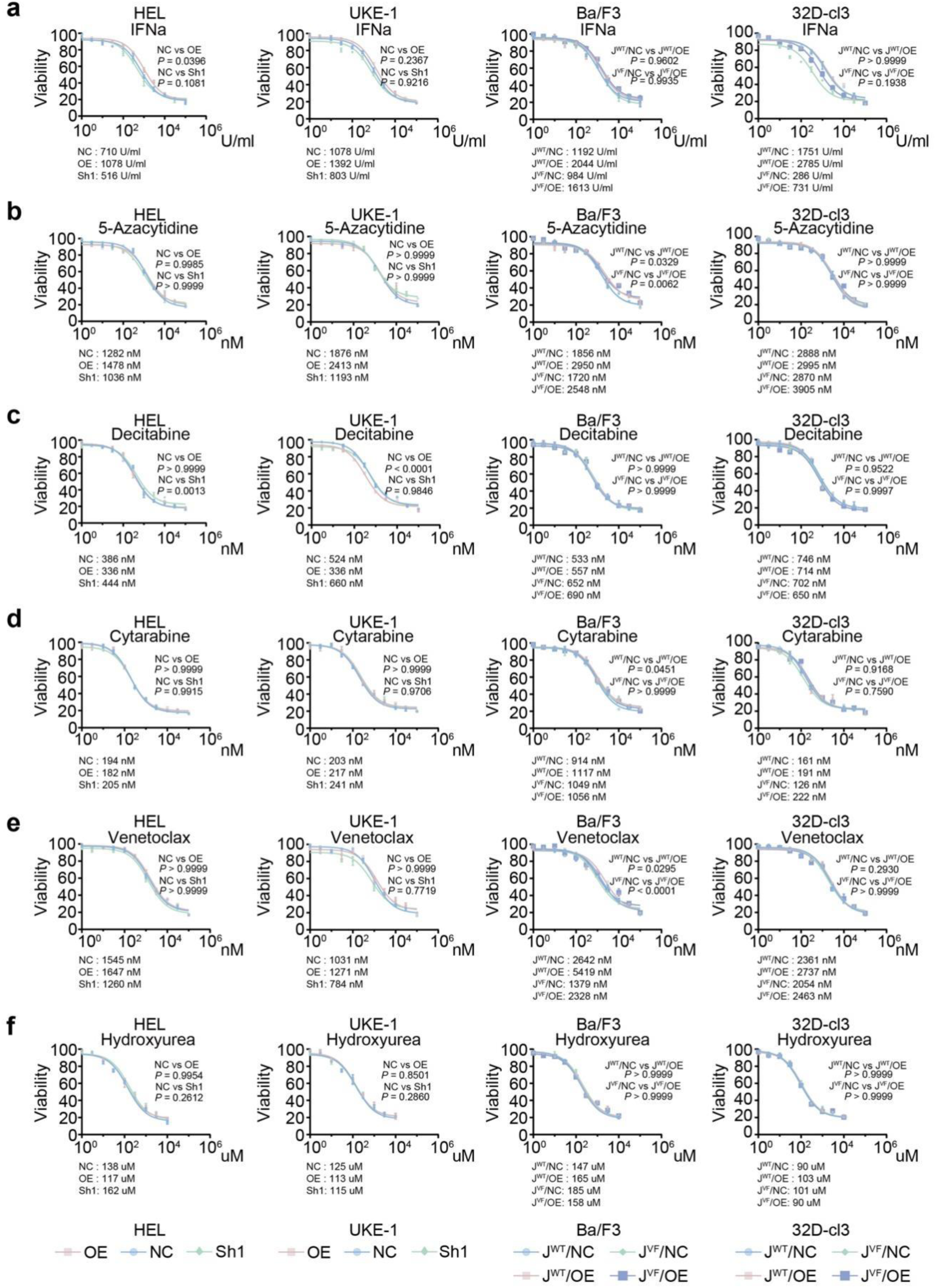
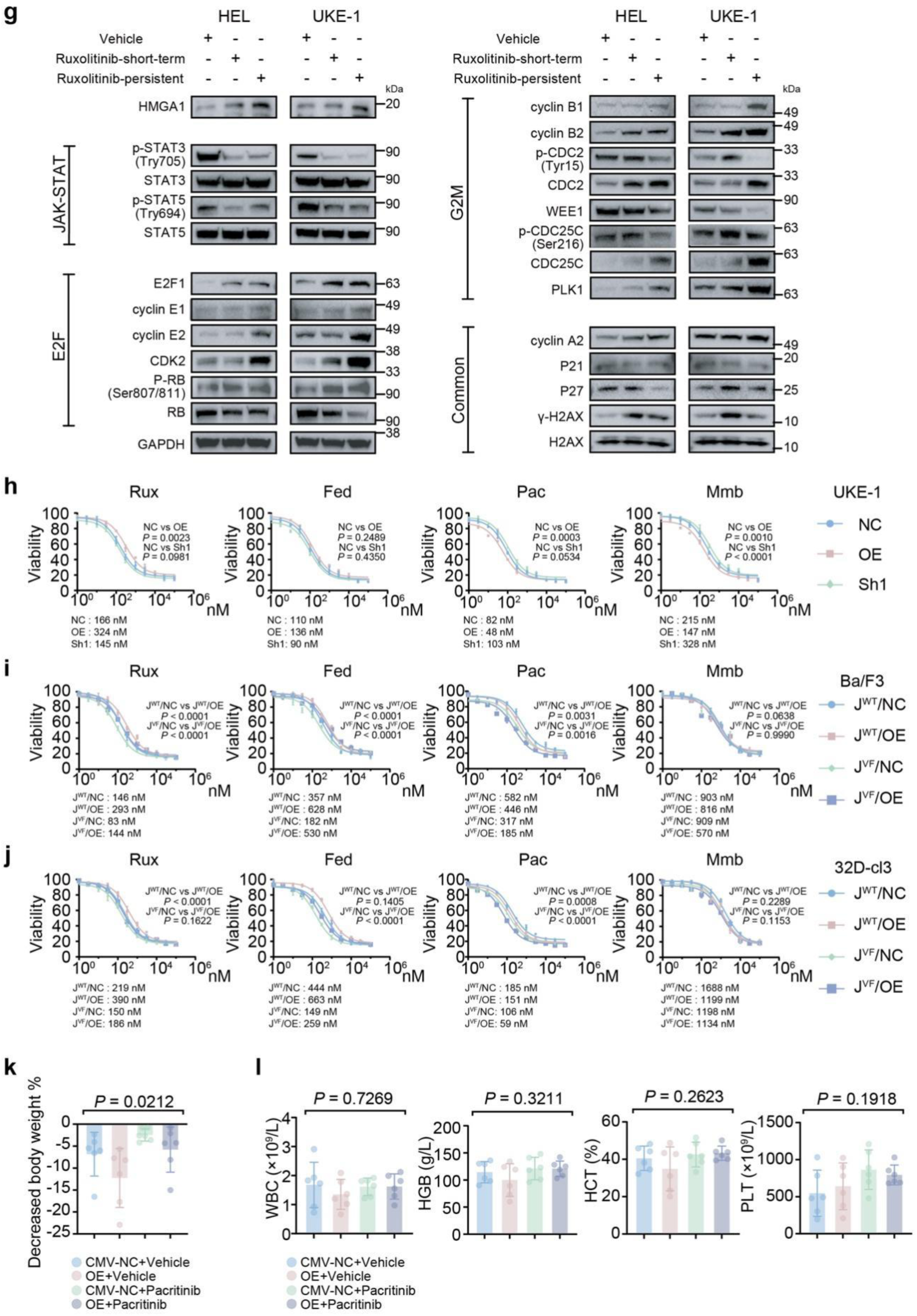
HMGA1 expression dictates therapeutic responses in MPN, and pacritinib demonstrates efficacy in HMGA1-driven disease. (a-f) HMGA1 expression levels modulate sensitivity to diverse therapeutic agents. Dose-response curves showing viability of HEL, UKE-1, Ba/F3 (*Jak2* wild type, or *Jak2*^V617F^), and 32D-cl3 (*Jak2* wild type, or *Jak2*^V617F^) cells with engineered HMGA1/Hmga1 expression (OE vs. NC; sh1/sh2 vs. sh-NC) following 72-hour treatment with (a) IFNα, (b) 5-Azacytidine, (c) Decitabine, (d) Cytarabine, (e) Venetoclax, and (f) Hydroxyurea. Calculated IC50 values are shown. Data represent mean ± SD from n = 3 independent experiments. Two-way ANOVA. (g) Ruxolitinib treatment, particularly long-term exposure, alters key signaling and cell cycle protein expression. Western blot analysis of indicated JAK-STAT, E2F pathway, G2M checkpoint, and cell cycle regulatory proteins in HEL and UKE-1 cells treated with vehicle, short-term ruxolitinib (4 hours), or in ruxolitinib-persistent (Rux-P) lines. GAPDH served as loading control. (h-j) HMGA1/Hmga1 expression status influences sensitivity to JAK inhibitors. Dose-response curves assessing viability of (h) UKE-1 cells, (i) Ba/F3 cells (J^VF^/NC: *Jak2*^V617F^/control vector; J^VF^/OE: *Jak2*^V617F^ /Hmga1 OE; J^WT^/NC: *Jak2* wild-type/control vector; J^WT^/OE: *Jak2* wild-type /Hmga1 OE), and (j) 32D-cl3 cells (similarly engineered) with engineered HMGA1/Hmga1 expression, following treatment with ruxolitinib, fedratinib, pacritinib, or momelotinib. Calculated IC50 values are shown. Data represent mean ± SD from n = 3 independent experiments. Two-way ANOVA. (k) Pacritinib mitigates weight loss in mice bearing HMGA1-overexpressing HEL xenografts. Percent body weight change in NSG mice engrafted with HEL-Luc cells (CMV-NC or HMGA1-OE) and treated with vehicle or pacritinib (100 mg/kg, BID, 14 days). Data are mean ± SD (n = 6 per group). One-way ANOVA. (l) Pacritinib treatment improves hematological parameters in the HMGA1-overexpressing HEL xenograft model. Peripheral blood counts (WBC, HGB, HCT, PLT) in xenografted mice at day 35 endpoint. Data are mean ± SD (n = 6 per group). One-way ANOVA.

